# CMDdemux: an efficient single cell demultiplexing method

**DOI:** 10.1101/2025.08.28.672983

**Authors:** Jianan Wang, Lizhong Chen, Daniel V. Brown, Chris Y. Chiu, Terence P. Speed

## Abstract

Multiplexing technologies label cells with molecular tags, allowing cells from different donors to be pooled together for sequencing. Although this approach enhances cell throughput, eliminates batch effects, and enables doublet detection, limitations of hashtag-based labelling can still lead to low-quality data. Existing demultiplexing methods can accurately assign donor identities in high-quality datasets, but they often fail on low-quality data. To address this, we developed CMDdemux, a method comprising three key steps: within-cell centered log-ratio (CLR) normalization of hashtag count data, K-medoids clustering, and classification of cells based on Mahalanobis distance. By integrating both hashing and mRNA data, CMDdemux achieves high accuracy in distinguishing singlets, doublets, and negatives. It also provides visualization tools to help users inspect potentially misclassified droplets. We benchmarked CMDdemux against existing methods using a range of high- and low-quality datasets. Results show that CMDdemux consistently outperforms other approaches, demonstrating robust performance on both high- and low-quality data where other methods fail. CMDdemux is particularly effective in handling diverse types of low-quality multiplexing data across different multiplexing technologies.

## Introduction

Single-cell technology has revolutionized the study of transcriptomics at single-cell resolution. The advancement of this technology requires the parallel sequencing of large numbers of cells. Although current sequencing platforms can capture profiles from tens of thousands of cells, they are typically limited to processing individual samples. Multiplexing—labelling each cell with a sample-specific barcode—addresses this limitation. Cell hashing[1] is the most widely used multiplexing method, which utilizes oligonucleotide-tagged antibodies targeting ubiquitously expressed surface proteins to label cells from different samples. MULTI-seq[2], another popular approach, achieves multiplexing through the hybridization of lipid- and cholesterol-modified oligonucleotides. Nucleus hashing[3] uses anti-nuclear pore complex antibodies to stain cells, making it suitable for complex tissues that are difficult to dissociate. Other advanced multiplexing strategies include the transient transfection of short barcoding oligonucleotides[4], labelling permeabilised nuclei treated with perturbations using single-stranded DNA[5], and incorporating short oligo barcodes into the green fluorescent protein (GFP) gene via lentiviral delivery[6]. These approaches enable multiplexing at various scales of throughput. Once labelled, cells are pooled for sequencing, and the process of assigning each cell back to its original sample by tracking the sample-specific barcodes is called demultiplexing. There are three demultiplexing categories: negatives, singlets, and doublets. Negatives are droplets that lack sufficient labelling signals; singlets are droplets with labelling signals primarily from a single donor; and doublets are droplets containing signals likely derived from multiple donors.

Multiplexing technologies offer several advantages for single-cell studies. First, they increase both sample and cell throughput. While previous experiments could process only one sample at a time, multiplexing enables the simultaneous sequencing of cells from multiple distinct samples. As a result, multiplexing also reduces costs and saves reagents and time. Additionally, multiplexing helps mitigate batch effects, as all cells are pooled and sequenced together rather than in separate runs. It also enhances the reliability of doublet detection. mRNA-based doublet detection methods[7–9] often rely on library size; however, transcriptomic profiles can vary widely in library size, and this variation is associated with cell type[10]. Consequently, these methods may fail to classify doublets accurately and risk discarding valid cells. In contrast, the additional modality provided by multiplexing—specifically, hashing signals introduced during the molecular experiment—offers a more robust and reliable means of demultiplexing. Another approach, SNP-based demultiplexing, determines sample identity using single-nucleotide polymorphisms (SNP) and includes methods such as demuxlet[11], Vireo[12], Souporcell[13], and scSplit[14]. These are typically considered accurate and are often used as ground truth in benchmarking studies. However, it is important to note that genotype-free methods are generally less accurate than genotype-based SNP demultiplexing, and in some cases perform worse than hashing-based methods. This may be due to the inherent difficulty in inferring SNPs from transcriptomic data and the reliance of such inference on mRNA quality.

While multiplexing technology offers many benefits, it also has limitations that can result in low-quality data. Issues such as improper staining[15], varying antibody sensitivity across specific cell types[16], non-standard experimental procedures, and sequencing errors can lead to problems in cell hashing data—namely, insufficient signals, contaminated signals, and abnormally strong signals. Fig. 1 provides an illustration of various types of multiplexed cell hashing data. In high-quality (standard) data, the signal distribution typically follows a bimodal pattern: a large peak with a small mean represents background noise, and a smaller peak with a larger mean represents true signals. This setting often results in three well-separated clusters upon visualization. Uneven labelling is a common issue in low-quality cell hashing. One such case involves weak signals, also referred to as unlabelled hashes. Here, one or more hashtags fail to bind tightly to cells, resulting in weak signals (low counts) and left-shifted distributions. These cells are often misclassified as negatives. Another form of uneven labelling is over-labelling, also known as strong signals. In this scenario, one or more hashes produce excessively strong signals across all cells, resulting in right-shifted distributions. The high counts of over-labelled hashes can cause demultiplexing algorithms to misclassify unaffected cells as doublets. Contamination is another typical type of low-quality cell hashing. It manifests as a group of cells labelled by multiple hashes, forming a contaminated cluster. This cluster shows moderate expression across all hashes and corresponds to the second peak in a tri-modal distribution—representing noise, contamination, and true signal peaks, respectively. Contaminated clusters usually consist of a mixture of negatives, singlets, and doublets. Cross-contamination often occurs during cell pooling[17], leading to hash labels from multiple donors attaching to the same cells. Another atypical low-quality type involves empty droplets—cells lacking any hash label. These droplets show low counts across all hashes and, if present in large numbers, may cluster together as negatives. Low-input data represent another challenge, characterized by low expression levels across all hashes. Their distributions often exhibit two closely spaced peaks, making it difficult to distinguish between noise and signal. Due to their similar expression profiles, cells in low-input data tend to group into indistinct clusters.

**Fig. 1.**
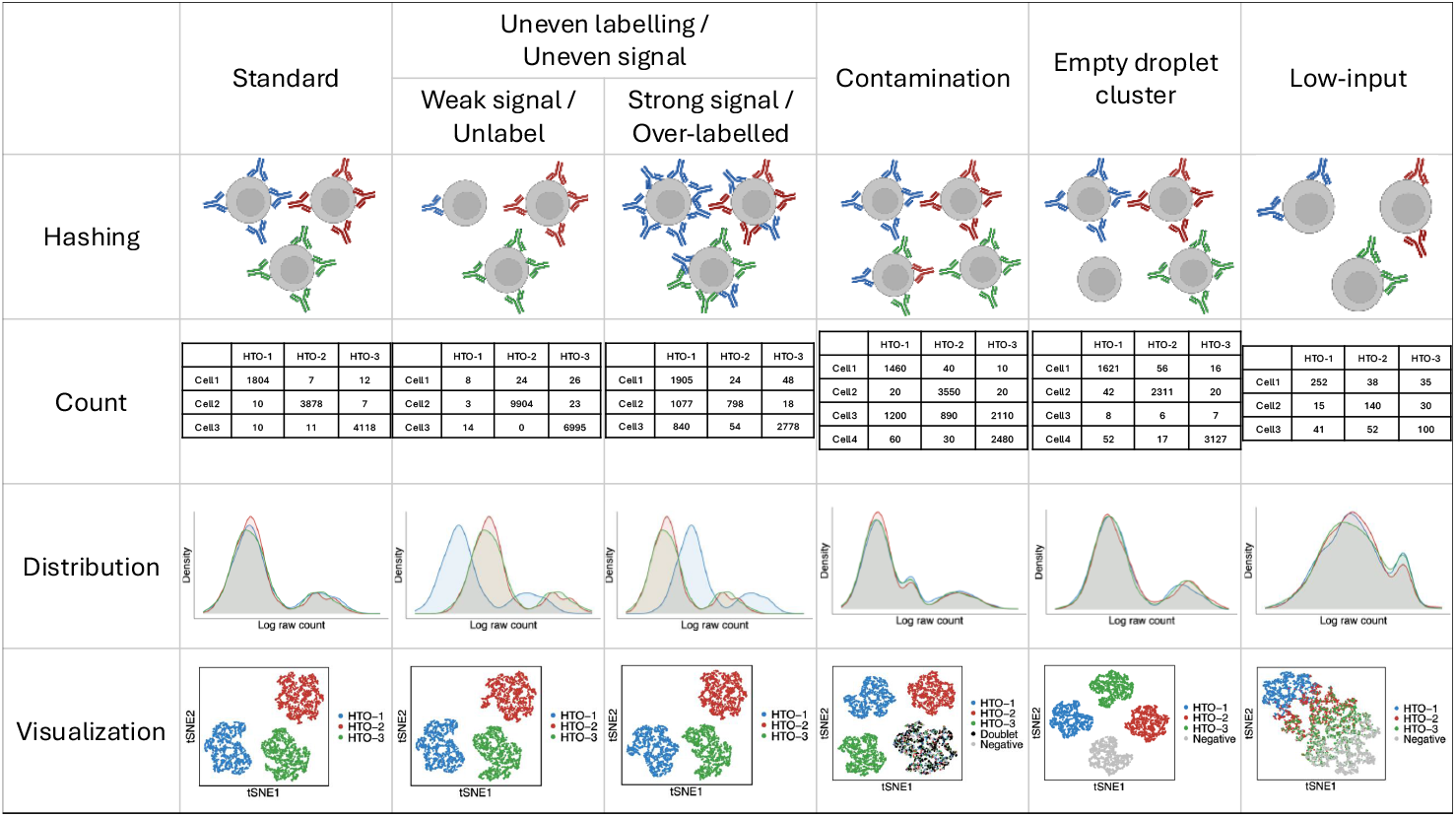
A summary of multiplexing hashing data types. An example involving three hashtag labels is shown in the figure. These hashtags do not follow a standard bimodal distribution, making it difficult to distinguish background noise from true signals and resulting in misclassification by demultiplexing algorithms. Empty droplets differ from weak signals in that weak signal data still display low-level signals from one or more hashtags, whereas empty droplets show no detectable signals from any specific hashtag. Low-input data are also distinct from weak signals: low-input refers to low expression across all hashtags, while weak signals involve low expression from only one or a few hashtags, with others remaining at standard levels. Contaminated data are different from over-labelled data. Contamination presents intermediate expression across multiple hashtags in a subset of cells, whereas over-labelled data show one or more hashtags with excessively high expression across all cells.

A variety of demultiplexing tools have been developed in recent years. A summary of these methods and their limitations is shown in Table S2. HTODemux[1] applies across-cell centered log-ratio (CLR) normalization—referred to here as global CLR normalization—followed by k-medoids clustering, fitting a negative binomial distribution to non-negative clusters, and setting a classification threshold based on the distribution’s quantile. hashedDrops[18] classifies cells according to the log fold change between the most abundant and the second most abundant hashtag. DemuxEM[3] uses an expectation-maximization (EM) algorithm to estimate the proportion of hashing counts from background noise and true signals. GMM-Demux[19] applies global CLR normalization, fits the normalized data to a two-component Gaussian mixture model, and classifies cells using Bayes’ posterior probabilities. BFF_raw_[20] fits the logtransformed hash counts to a kernel density curve and sets the threshold at the point of lowest density. BFF_cluster_[20] normalizes hashtag oligonucleotide (HTO) counts using bimodal quantile normalization (BQN), which separately normalizes positive and negative cells, combines the normalized values, and applies a threshold to classify cells. demuxmix[21] fits a regression model between HTO counts and the number of detected genes from the mRNA profile. deMULTIplex2[22] uses a negative binomial generalized linear model (GLM-NB) between contaminated HTO counts and total HTO counts. Despite their strengths, these algorithms share some common limitations. Unsurprisingly, all methods perform well on high-quality data, as they were developed and tuned using such datasets. However, none of them performs consistently well on all types of low-quality data. This is primarily because low-quality data often do not follow a standard bimodal distribution, violating model assumptions. Another contributing factor is the use of default cutoffs—optimized for high-quality data—that do not generalize well to low-quality cases. Normalization strategy also plays a critical role. Methods like global CLR and BQN aim to unify the distribution of hashtags, which works well when hashtags are similarly distributed (as in high-quality data). However, low-quality data often display different signal distributions, and inappropriate normalization can lead to misclassification. Therefore, the choice of normalization method significantly affects the performance of demultiplexing tools, especially on low-quality data.

Here, we introduce a novel demultiplexing method, CMDdemux, which uses within-cell CLR normalization and Mahalanobis distance[23] to assign cells to their donor of origin. We first perform CLR normalization across hashtags within each cell. This local CLR normalization strategy preserves the characteristics of each hashtag while adjusting for within-cell variation. Next, cells are clustered in the local CLR space using a k-medoids clustering approach. To reduce the influence of outliers, only core cells—those close to the centroid of each cluster—are used to compute the covariance matrix. Mahalanobis distances are then calculated based on the cluster centroid, the local CLR values, and the cluster-specific covariance matrix. Unlike many existing methods, CMDdemux does not assume a bimodal distribution of hashtag expression, which allows it to deliver robust and accurate performance on both high-quality and low-quality data. Benchmarking results across multiple datasets show that CMDdemux performs especially well on diverse types of low-quality data generated by various multiplexing techniques.

## Results

### An overview of CMDdemux

The CMDdemux workflow includes three major steps: (1) within-cell CLR normalization, (2) K-medoids clustering, and (3) Mahalanobis distance-based classification (Fig. 2).

**Fig. 2.**
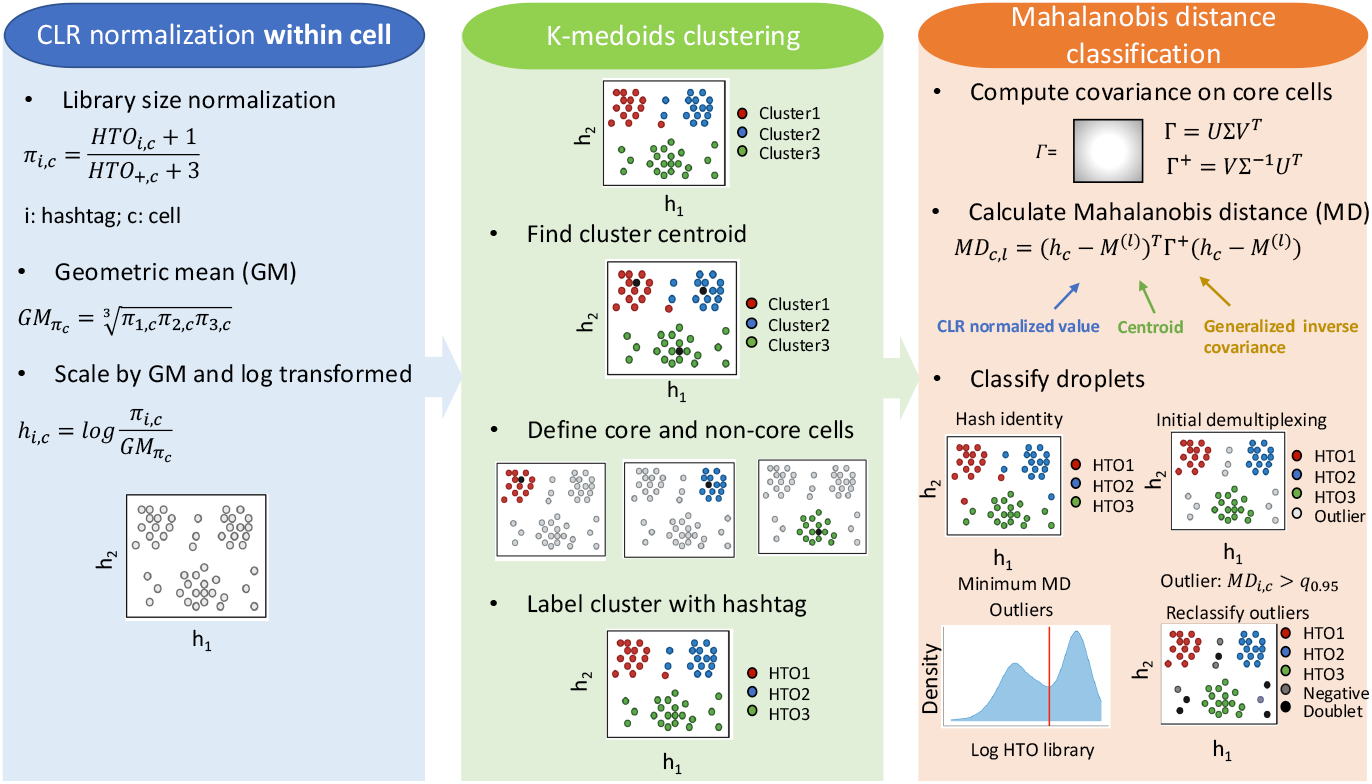
Schematic workflow of CMDdemux. The figure illustrates an example using three hashtags. CMDdemux first performs CLR normalization within each cell, referred to as local CLR normalization in this paper. K-medoids clustering is then applied to the local CLR normalized values. Finally, Mahalanobis distances are calculated using the outputs from the previous two steps. Demultiplexing is primarily based on the minimum Mahalanobis distance. A pseudo-count is added to each entry in the hash count matrix to avoid issues with log transformation of zero values.

In contrast to the across-cell CLR normalization strategy used by HTODemux and GMM-Demux, we refer to the CLR normalization performed within each cell as local CLR normalization (see “Methods”), and the normalization across cells as global CLR normalization (see Supplementary Methods Section S1). For local CLR normalization, library size normalization is first applied, followed by scaling by the geometric mean across all hashtags. The logarithm of this scaled value yields the local CLR normalized value. These local CLR values are used in all subsequent steps and for visualization.

K-medoids clustering is then performed on the local CLR normalized values. Cluster centroids are identified, and cells are classified as either core or non-core based on their Euclidean distance to the centroid. Core cells are those closer to the cluster centroid, exhibiting smaller Euclidean distances, while non-core cells are located near the cluster edges and have larger Euclidean distances. Defining core and non-core cells helps identify potential outliers within clusters, which may affect the within-cluster and between-cluster variance in subsequent steps. Finally, clusters are assigned back to their sample of origin.

A covariance matrix of the clusters is computed using only the core cells. To avoid issues with matrix singularity, a generalized inverse of the covariance matrix is used. The Mahalanobis distance (MD) is then calculated based on the local CLR normalized values, cluster centroids, and the covariance matrix. Each droplet is classified to the hashtag with the minimum MD. In the initial demultiplexing step, droplets with relatively large minimum MD values are defined as outliers. These outlier cells are likely to be either negatives or doublets, and their hashtag library sizes are expected to have a bimodal distribution—one mode representing negatives (lower expression) and the other representing doublets (higher expression). The antimode of this distribution is used as a threshold to further reclassify outliers into negatives and doublets.

### CMDdemux provides a visual check for suspicious droplets

Compared with other methods, one advantage of CMDdemux is that it provides a visual check for suspicious droplets. The functions CheckAssign2DPlot and CheckAs-sign3DPlot generate plots that allow users to inspect potentially misclassified droplets. The CheckAssign2DPlot function displays pairwise relationships between log HTO library size, log mRNA library size, second Mahalanobis distance, and the ratio of the first to second Mahalanobis distances in two-dimensional (2D) plots (Fig. 3a). The CheckAssign3DPlot function visualizes the log HTO library size, log mRNA library size, and the ratio of the first to second Mahalanobis distances along the three axes of a three-dimensional (3D) plot (Fig. 3b). In both 2D and 3D plots, singlets, doublets, and negatives are separated into distinct regions. Users can specify cell barcodes of interest, and their locations in the plots help diagnose potential misclassifications. This provides a quick and effective way to evaluate demultiplexing results, particularly when SNP-based demultiplexing is not available.

**Fig. 3.**
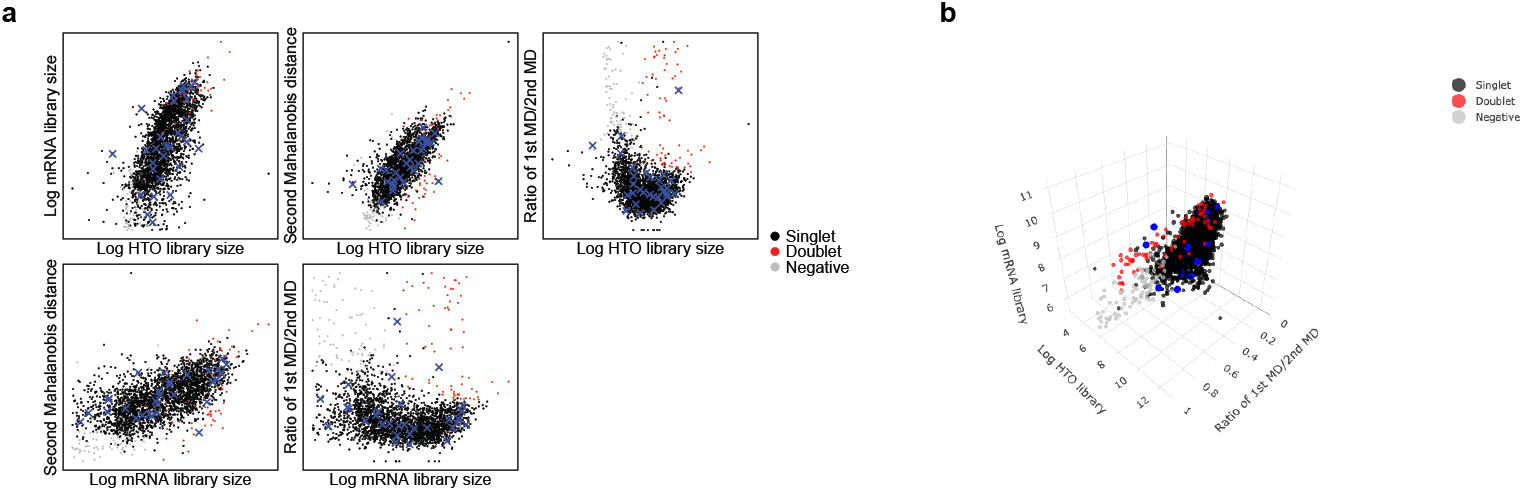
Demonstrate visual check by CMDdemux. The demonstrated plots are based on a high-quality human brain data. Thirty singlets are randomly selected and are labelled as blue dots in the plots. (a) Plots are generated by CheckAssign2DPlot function. (b) The plot is generated by CheckAssign3DPlot function.

### CMDdemux outperforms other methods on data with weak signals

In low-quality datasets, it is quite common for some hashtags not to bind tightly to cells. These loosely bound hashtags result in weak signals, which appear as low HTO counts for certain unlabelled hashtags. Such weak signals disrupt the expected pattern distinguishing background noise from true hashtag signals. As a result, many demultiplexing algorithms may produce misleading assignments, failing to correctly classify cells that should be associated with those hashtags. In this section, we present two example datasets with weak signals and no ground truth.

The mouse dataset[24] provides an example with an unlabelled hashtag — specifically, mouse2. It is evident that the mouse2 hashtag does not bind tightly to the cells. Compared to the other two hashtags, mouse2 shows lower HTO library sizes (Fig. 4a). In addition, the distribution of the mouse2 hashtag is non-bimodal and left-shifted, whereas the other two hashtags exhibit similar bimodal distributions (Fig. S3a). Clusters 1, 2, and 3 can be reasonably assigned to mouse1, mouse2, and mouse3, respectively (Fig. S3b). Among the tested methods, CMDdemux is the only one that successfully demultiplexes both mouse1 and mouse2 cells. In contrast, HTODemux, GMM-demux, and demuxmix misclassify mouse1 cells as doublets and only partially identify mouse2 singlets (Fig. 4b). BFF cluster also misassigns the mouse2 cluster as doublets. These misassignments occur because the mouse2 hashtag shows similarly low expression levels in both the mouse1 and mouse2 clusters (Fig. S3c), leading to confusion between true singlets and doublets. Methods like demuxEM, hashedDrops, and BFF raw classify cells in the mouse2 cluster as negatives due to the weak mouse2 signal. Lastly, deMULTIplex2 fails to produce an output for this dataset, likely because the small counts do not fit the negative binomial generalized linear model (GLM-NB) well.

**Fig. 4.**
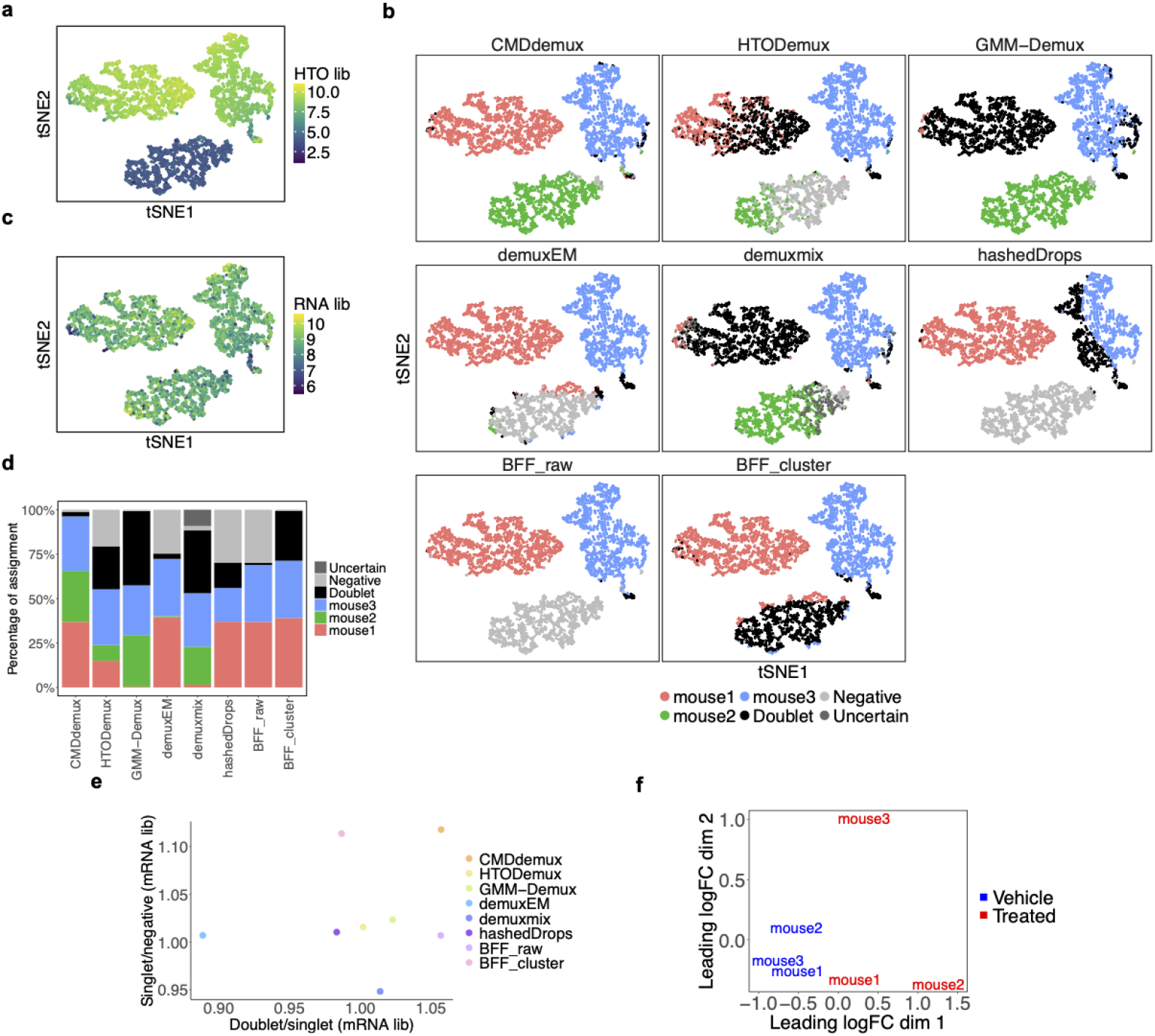
Performance of CMDdemux on the treated mouse dataset. (a–c) t-SNE plots based on local CLR-normalized HTO values. (a) Colored by HTO library sizes. (b) Colored by demultiplexing results from different methods. deMULTIplex2 fails to produce demultiplexing results. (c) Colored by mRNA library sizes. (d) Proportion of cell assignments by different methods. (e) Median mRNA library sizes for each demultiplexing category are shown. The x-axis represents the ratio of the median mRNA library size in doublets to that of the singlet group with the highest median mRNA library size. The y-axis represents the ratio of the median mRNA library size of the singlet group with the lowest value to that of the negative group. An ideal demultiplexing method should show high values on both axes. (f) Multidimensional scaling (MDS) plot of pseudo-bulk samples based on RNA expression profiles. Pseudo-bulk samples were constructed using CMDdemux demultiplexing labels.

Upon further examination of the mRNA library sizes, the three clusters fall within a reasonable range (Fig. 4c), supporting their assignment as singlets rather than as clusters of doublets or negatives. In most experiments, cells from different samples are expected to be labelled evenly, yielding a similar proportion of cells across samples. CMDdemux is the only method that provides a balanced singlet assignment across the three hashtags (Fig. 4d). Compared to other methods, CMDdemux also assigns a reasonable proportion of doublets and negatives (Fig. S3d). The doublets identified by CMDdemux tend to have higher mRNA library sizes than singlets, while negatives show lower mRNA library sizes. Additionally, singlets from all three samples exhibit similar mRNA library sizes (Fig. S3e). Among all methods, CMDdemux yields the highest ratio of mRNA library size in doublets to singlets, compared to the ratio of singlets to negatives, reinforcing the reliability of its assignments (Fig. 4e). To further validate the demultiplexing results, we computed the silhouette scores. CMDdemux achieves high scores for mouse1 and mouse3 hashtags and a moderate score for mouse2 (Fig. S3f). The median silhouette score across all singlets is also high, indicating strong clustering performance (Fig. S3g).

Another example dataset is the EMBRYO MULTI-Seq LMO data[25]. In this dataset, it is clear that MULTI 2 labels only a few cells (Fig. 5a). Because the signal of MULTI 2 is too weak, the cells labelled by MULTI 2 are scattered rather than forming a distinct cluster. Moreover, the distribution of the MULTI 2 hash differs sub-stantially from other hashtags, with most cells showing very weak signals (Fig. S4a). This makes it challenging for algorithms to accurately detect cells from the MULTI 2 sample. CMDdemux implements a specific method to identify cells from such weakly labelled hashtags, as described in Supplementary Methods Section S4.

**Fig. 5.**
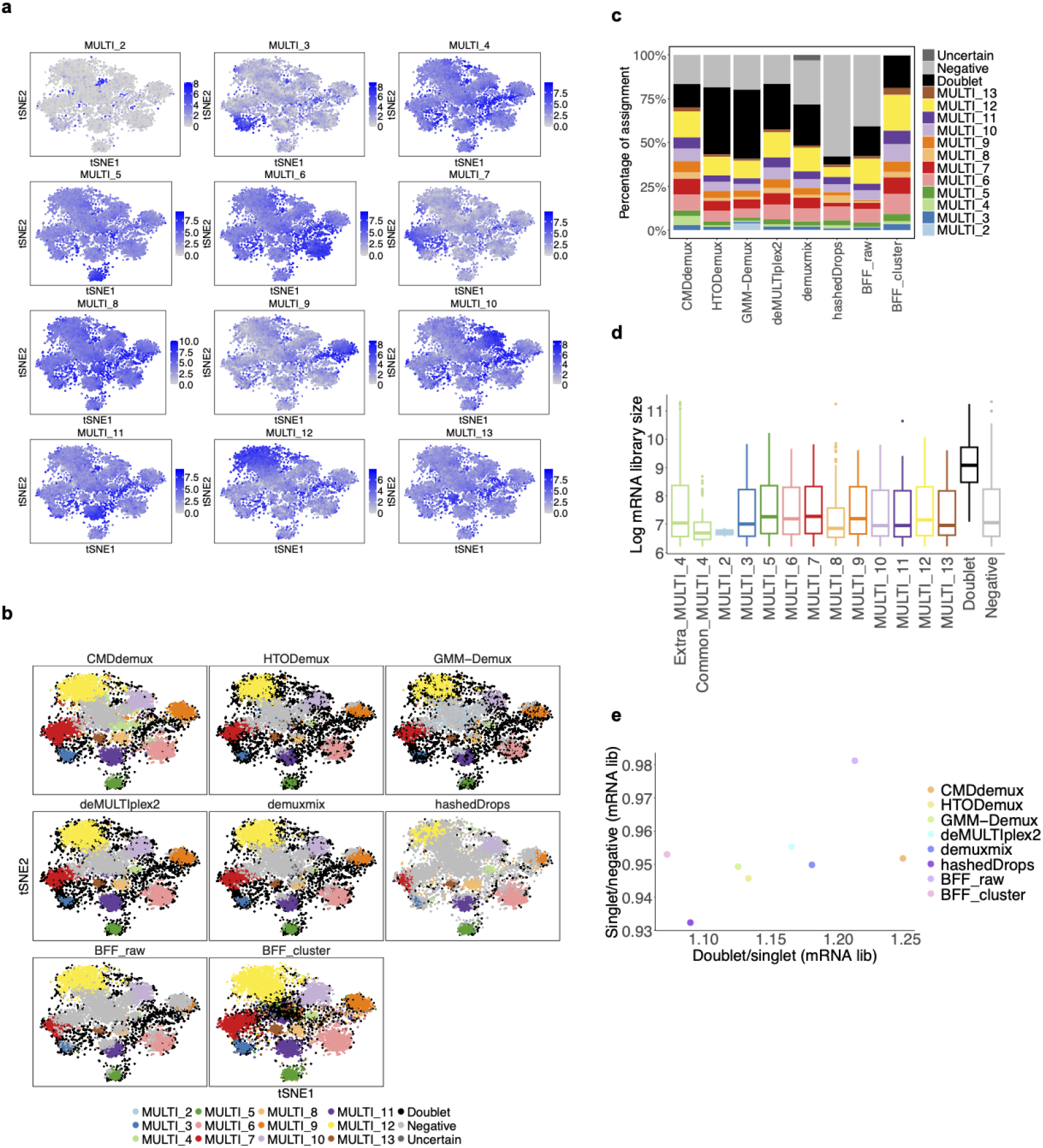
Performance of CMDdemux on the EMBRYO LMO data. (a) Each t-SNE plot is colored by the log count of the corresponding hashtag. (b) t-SNE plots colored by demultiplexing results from different methods. demuxEM is unable to generate demultiplexing results. (c) Proportion of assignments by different methods. (d) Distribution of log mRNA library sizes for different demultiplexing categories as determined by CMDdemux. “Common MULTI 4” refers to MULTI 4 singlets assigned by CMDdemux, deMULTIplex2, hashedDrops, and BFF cluster. “Extra MULTI 4” refers to additional MULTI 4 singlets assigned only by CMDdemux. (e) The horizontal axis represents the ratio of the median mRNA library size in doublets to that in the singlet category with the highest median mRNA library size. The vertical axis represents the ratio of the median mRNA library size in the singlet category with the lowest median to that in all negatives.

The main difference between CMDdemux and other methods is that CMDdemux assigns more MULTI 4 singlets compared to the others (Fig. 5b, c). Some singlets in each singlet cluster are not detected by demuxmix, hashedDrops, and BFF raw. These three methods assign fewer singlets and classify those cells as negatives, likely due to their stringent thresholds for distinguishing singlets from negatives. The BFF cluster method assigns droplets in the middle cluster as doublets, whereas all other methods classify that cluster of cells as negatives. However, that cluster does not exhibit elevated HTO library sizes (Fig. S4b) or higher mRNA library sizes (Fig. S4c), so it should not be classified as doublets. HTODemux and GMM-Demux assign more doublets (Fig. S4d) than other methods, mostly at the edges of the singlet clusters (Fig. 5b). Nevertheless, these cells have reasonable HTO (Fig. S4b) and mRNA library sizes (Fig. S4c). Cells located between clusters that exhibit higher HTO and mRNA library sizes are more likely to be true doublets.

We further examined the additional MULTI 4 singlets assigned by CMDdemux. These cells show mRNA library sizes comparable to other singlet categories, and are similar to the common MULTI 4 singlets assigned by other methods (Fig. 5d). In addition, randomly selected extra MULTI 4 singlets tend to be located in regions corresponding to singlets, rather than negatives or doublets (Fig. S4e), further supporting the validity of CMDdemux’s assignments. Overall, CMDdemux identifies doublets with higher mRNA library sizes (Fig. S4f). It also yields higher ratios of mRNA library sizes between doublets and singlets, and between singlets and negatives (Fig. 5e, S4g). A high Calinski–Harabasz (CH) index further indicates CMDdemux’s strong performance in singlet demultiplexing (Fig. S4h).

Low-quality cell hashing data with weak signals can result from multiple factors. One important factor is the cell state. Treated mice exhibit different cellular states compared to vehicle-treated mice (Fig. 4f). Drug treatment can alter cell states, which in turn affects the binding efficiency of hashing antibodies to the cells. Moreover, since the mouse hashtag antibodies target CD45 and MHC class I, and the cells are CD45-sorted by flow cytometry, drug treatment may influence the expression levels of these proteins, thereby affecting the binding efficiency of the hashtag antibodies. In the case of the EMBRYO MULTI-Seq LMO data, according to the original paper, the unlabelled hash may have been caused by incomplete removal of supernatants. The lipid hashtags from MULTI-Seq are quenched by serum in the wash buffer. If the wash buffer is not properly removed, residual proteins in the medium may interfere with the binding efficiency of the hashtag oligos.

### CMDdemux demonstrates superior performance on the data with excessively strong signals

In contrast to the condition where some hashtags show weak signals, there is another scenario in which certain hashtags exhibit more intense signals compared to those that typically bind to cells. Generally, hashtags with excessively strong signals display a right-shifted distribution in the hashing count. This right-shift causes the true signals of normally labelled hashtags to overlap with the background noise of the over-labelled hashtags. As a result, some cells may show high counts for both normally labelled and over-labelled hashtags. However, the high counts from the over-labelled hashtags likely represent noise due to their over-labelling nature. These elevated counts from multiple hashtags may mislead some algorithms into incorrectly classifying those cells as doublets.

The OT data[26] is a typical example of a dataset with an over-labelled hashtag. The distribution of OT-G is right-shifted compared to the normally labelled hashtags OT-E and OT-F (Fig. 6a), and the noise from OT-G overlaps with the signal ranges of other hashtags. Due to over-labelling, OT-G signals are more intense than those from other hashtags (Fig. S5a). As a result, nearly all cells exhibit higher counts of OT-G. Consequently, it is not surprising that demuxEM and BFF cluster report an unreasonably large number of doublets (Fig. 6b). Other methods, such as HTODe-mux, GMM-Demux, demuxmix, hashedDrops, and BFF raw, also struggle to correctly classify OT-G singlets, often misclassifying them as negatives. One key reason is that OT-G exhibits a unimodal distribution (Fig. S5b), which violates the assumptions of most methods—namely, that hashtags follow a bimodal distribution with one peak for noise and another for signal. This unimodal distribution complicates the determination of a clear signal threshold.

**Fig. 6.**
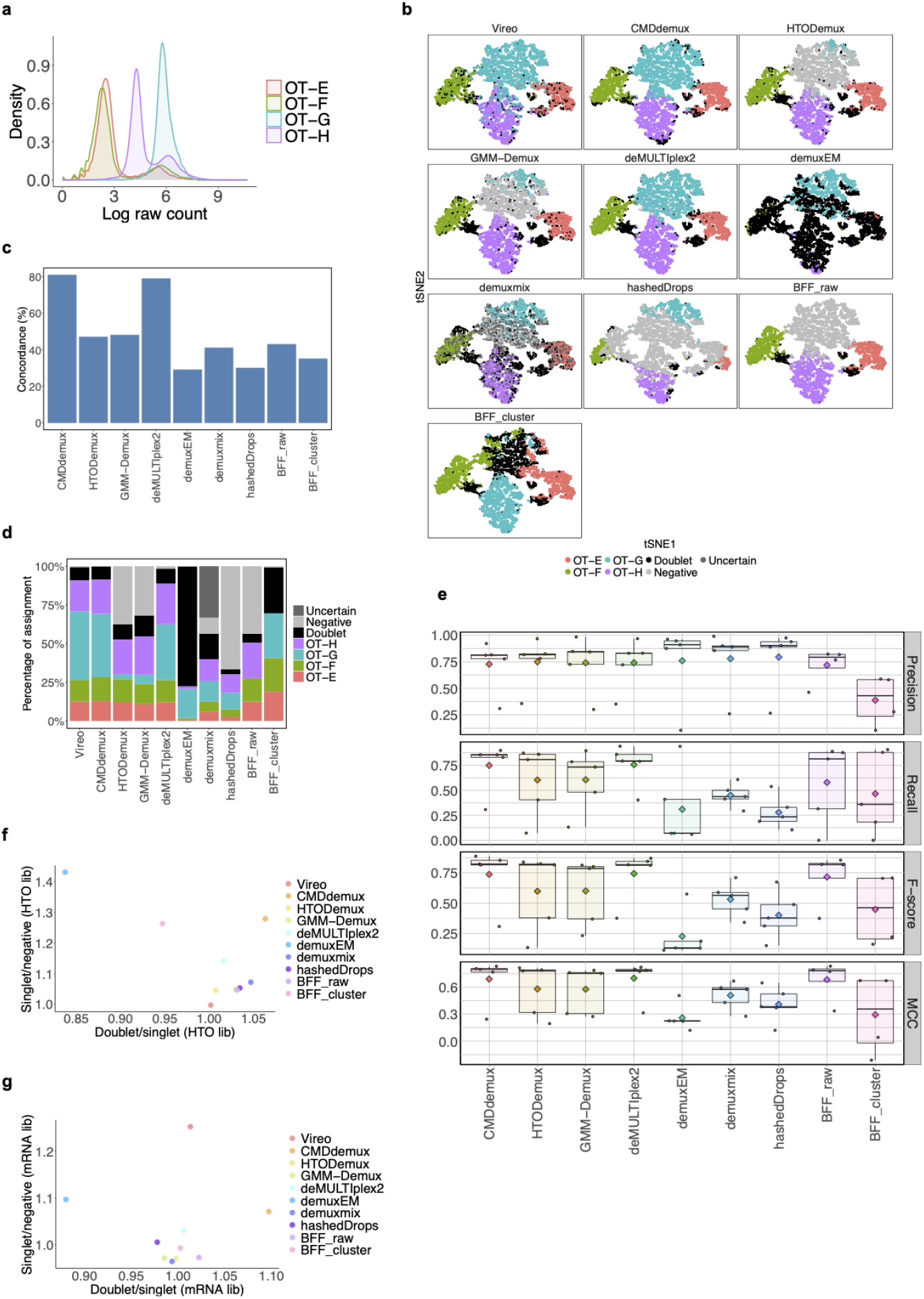
Performance of CMDdemux on the OT data. We treat the Vireo-derived assignments as the ground truth for benchmarking purposes. (a) Distribution of log-transformed raw hashtag count data across all cells for each hashtag. (b) t-SNE plot colored by demultiplexing results from different methods. (c) Total concordance with Vireo, calculated as the number of cells classified identically to Vireo divided by the total number of cells in the OT dataset. (d) Proportion of cell assignments to different categories by each method. (e) Performance evaluation of each method using precision, recall, F1 score, and Matthews correlation coefficient (MCC). Each dot represents a demultiplexing category (i.e., each singlet and doublets); negatives are excluded. Diamonds indicate mean values. (f, g) The x-axis represents the ratio of median library sizes between doublets and singlets (the singlet category with the highest median), and the y-axis represents the ratio between singlets (the singlet category with the lowest median) and negatives. (f) HTO library sizes. (g) mRNA library sizes.

Using the SNP-based demultiplexing method Vireo as the ground truth, both CMDdemux and deMULTIplex2 produced similar demultiplexing results (Fig. 6b). These two methods showed high concordance with Vireo’s classifications (Fig. 6c, S5c), with CMDdemux achieving 81% concordance and deMULTIplex2 achieving 79%. CMDdemux also exhibited a classification distribution more closely aligned with Vireo’s (Fig. 6d). The performance of each method was further evaluated using precision, recall, F1 score, and Matthews correlation coefficient (MCC) (Fig. 6e, S5d). Both CMDdemux and deMULTIplex2 outperformed other methods in classifying singlets and doublets. Additionally, the high ratio of HTO library sizes—doublets to singlets and singlets to negatives (Fig. 6f)—as well as the corresponding mRNA library size ratios (Fig. 6g), suggest that CMDdemux produces reliable demultiplexing results. The mRNA library sizes assigned by CMDdemux also show clear distinctions among negatives, singlets, and doublets (Fig. S5e).

The PDX CellPlex dataset[25] also suffers from an over-labelling problem. It is evident that the hashtag CMO301 is right-shifted in distribution (Fig. 7a) and is highly expressed in the majority of cells, in contrast to other hashtags that are uniquely highly expressed in specific clusters (Fig. S6a). The high expression of CMO301 leads to elevated counts in most cells, making it difficult for some methods to correctly identify CMO301 singlets. Both deMULTIplex2 and BFF fail to produce reasonable demultiplexing results (Fig. 7b), particularly in accurately identifying CMO301 singlets. Other methods successfully demultiplex singlets for each sample. HTODemux and demuxmix identify fewer CMO301 singlets than expected. hashedDrops assigns fewer singlets per sample, while GMM-Demux overassigns doublets. CMDdemux assigns a reasonable proportion of singlets across samples (Fig. 7c), and its doublet rate and negative rate (Fig. S6b) are also acceptable.

**Fig. 7.**
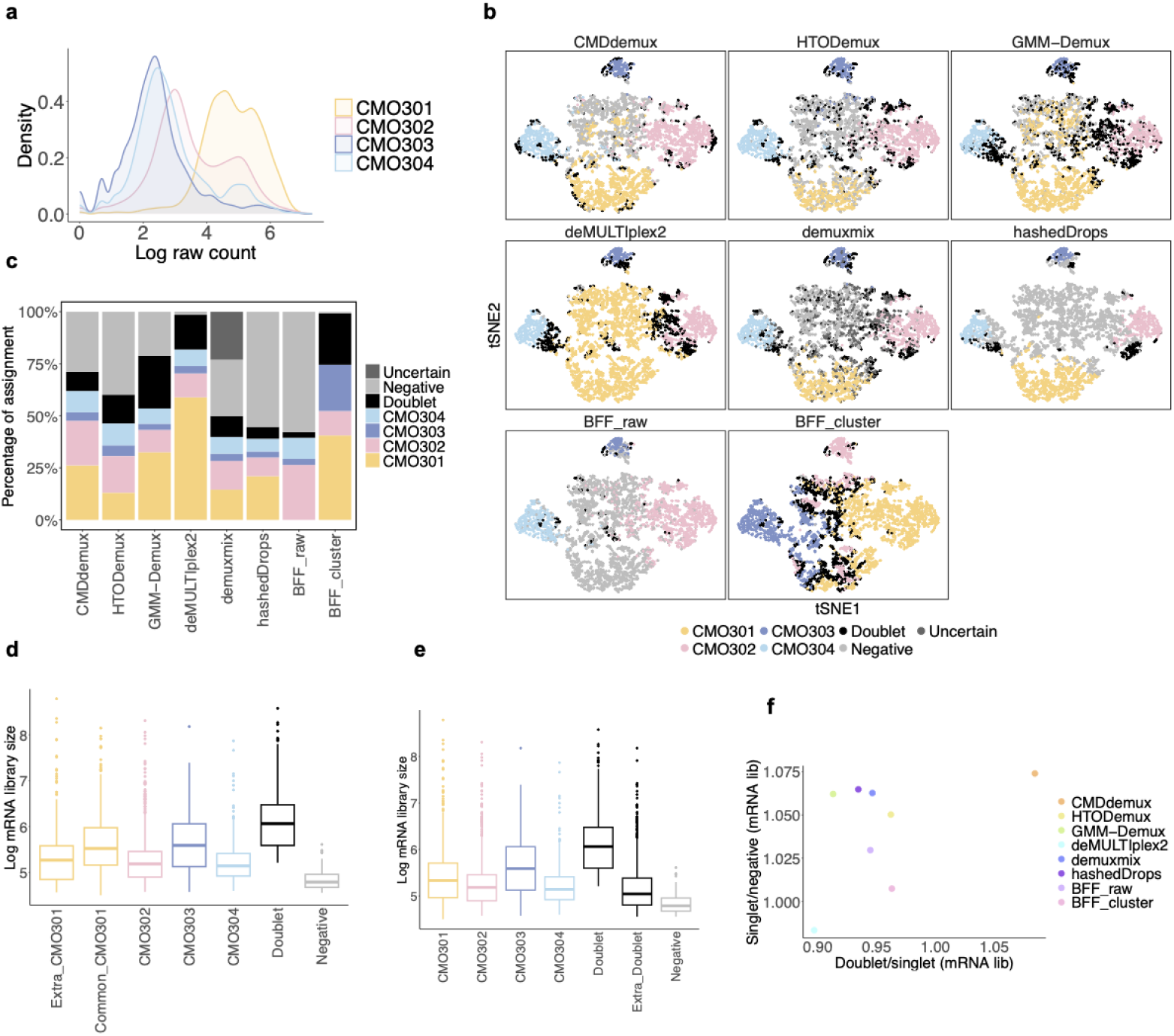
Performance of CMDdemux on PDX CellPlex data. (a) Distribution of raw counts for each hashtag. (b) t-SNE plot colored by demultiplexing results from different methods. demuxEM does not produce demultiplexing results. (c) Percentage of cells assigned to each category by different methods. (d, e) Boxplots of mRNA library sizes for each demultiplexing category. (d) “Common CMO301” refers to singlets identified by both HTODemux and demuxmix. “Extra CMO301” refers to singlets identified by CMDdemux but not by HTODemux or demuxmix. (e) “Extra Doublet” refers to doublets identified by GMM-Demux but not by CMDdemux. (f) The horizontal axis shows the median mRNA library size ratio between doublets and singlets (for the singlet category with the largest median mRNA library size). The vertical axis shows the median mRNA library size ratio between singlets (for the singlet category with the smallest median mRNA library size) and negatives.

We then further examined the additional CMO301 singlets assigned by CMDdemux and the extra doublets assigned by GMM-Demux. The additional singlets identified by CMDdemux have similar mRNA library sizes to singlets from other samples (Fig. 7d), and most of them are located within the singlet region (Fig. S6c). In contrast, the extra doublets identified by GMM-Demux have smaller mRNA library sizes (Fig. 7e), and most are also located in the singlet region (Fig. S6c). These findings further support that CMDdemux correctly assigns these cells as CMO301 singlets, whereas GMM-Demux misclassifies singlets as doublets. Moreover, compared to other methods, CMDdemux demonstrates a clearer separation of mRNA library sizes among negatives, singlets, and doublets (Fig. S6d). It also shows higher mRNA library size ratios between doublets and singlets, as well as between singlets and negatives (Fig. 7f, S6e).

Strong signals in specific hashtags may be caused by cell type differences. The cells in both the OT data and the PDX CellPlex data are tumour cells from ovarian carcinoma patients. Tumour cell states differ from those of normal cells, and some patients may have more advanced tumours than others. Additionally, this type of ovarian cancer is more difficult to sequence, as small metastases often lead to longer surgeries, increasing the risk of tissue degradation[27]. Differences in tissue pre-processing further contribute to inter-patient heterogeneity, which can ultimately result in varying degrees of adhesion to hashtag antibodies. For the PDX CellPlex data, nuclei were extracted to preserve tumor epithelial cells. Because nuclei are more fragile than whole cells, the additional processing required for nuclear preparation may introduce extra noise. Moreover, this preparation can generate more debris, likely from membrane fragments, which may bind non-specifically to CellPlex reagents. As a result, the low data quality may be due to microfluidic blockages caused by this debris, which can interfere with the even distribution of reagents across samples and lead to imbalanced signal-to-background ratios. This can be mitigated by determining the optimal concentration for specific binding, as well as by using Fc (fragment crystallizable) receptor blockade or having enough serum protein to quench hashtag antibodies and reduce non-specific binding.

### CMDdemux shows strong abilities in dealing with contaminated hashing data

When multiple hashtags are used to label cells, it is common for some hashtags to spill over into cells that were not originally intended to be labelled during the pooling process. When this happens, a cell may be labelled by multiple hashtags at approximately equal levels. This cross-contamination can lead some algorithms to misclassify singlets as doublets. Contaminated hashing data differ from data with excessively strong signals. In datasets with strong signals, one or more hashtags are overexpressed, causing cells labelled by other hashtags to also show high expression of the overexpressed tags. This can lead to misclassification or ambiguous labelling. In contrast, contaminated data typically contain a group of cells that express most hashtags at intermediate levels. These cells cluster together, forming a distinct contaminated cluster, while the remaining cells are normally labelled and show expected expression patterns for their assigned hashtags. The EMBRYO CMO dataset[25] is a typical example of such contamination. Each hashtag is highly expressed in a specific cluster but also moderately expressed in a larger group of cells (Fig. 8a) —referred to here as the contaminated cluster (cluster 5 in Fig. S7a). Unlike the classic bimodal distribution, each hashtag exhibits an approximately trimodal distribution (Fig. S7b), with peaks corresponding to background noise, contaminated signals, and true signals. This pattern is also reflected in the right-shifted hash count distribution in the contaminated cluster (Fig. 8b), where counts are higher than background noise but lower than true signal levels. Different methods show high concordance in demultiplexing cells from most clusters, except for the contaminated cluster (Fig. 8c). Notably, deMULTIplex2 assigns the majority of cells in the contaminated cluster as Nxt 460 singlets, which appears to be implausible. BFF raw and hashedDrops classify the entire contaminated cluster as negatives. However, the mRNA library sizes of cells in this cluster are similar to those in other singlet clusters (Fig. 8d, S7c). Therefore, the demultiplexing results from CMDdemux, HTODemux, GMM-Demux, demuxmix, and BFF cluster seem more plausible, as they suggest that the contaminated cluster likely consists of a mixture of negatives, singlets, and doublets. We further examined the CMDdemux-assigned cells in the contaminated cluster. These cells exhibit mRNA library sizes that are higher than those of negatives but lower than those of doublets, and their library sizes are comparable to those of the same demultiplexing category in other clusters (Fig. 8e). Additionally, using the CMDdemux examine function, randomly selected droplets from the contaminated cluster are correctly positioned in the areas corresponding to negatives, singlets, and doublets (Fig. S7d). Overall, both CMDdemux and BFF cluster provide a reasonable demultiplexing distribution across samples (Fig. 8f). CMDdemux also achieves a low negative rate and a realistic doublet rate (Fig. S7e). The CMDdemux-classified singlets in the contaminated cluster can help recover more usable cells for downstream analyses. CMDdemux demonstrates a clear separation of mRNA library sizes among negatives, singlets, and doublets (Fig. S7f). Moreover, it yields high ratios of median mRNA library sizes between doublets and singlets, as well as between singlets and negatives (Fig. 8g, S7g).

**Fig. 8.**
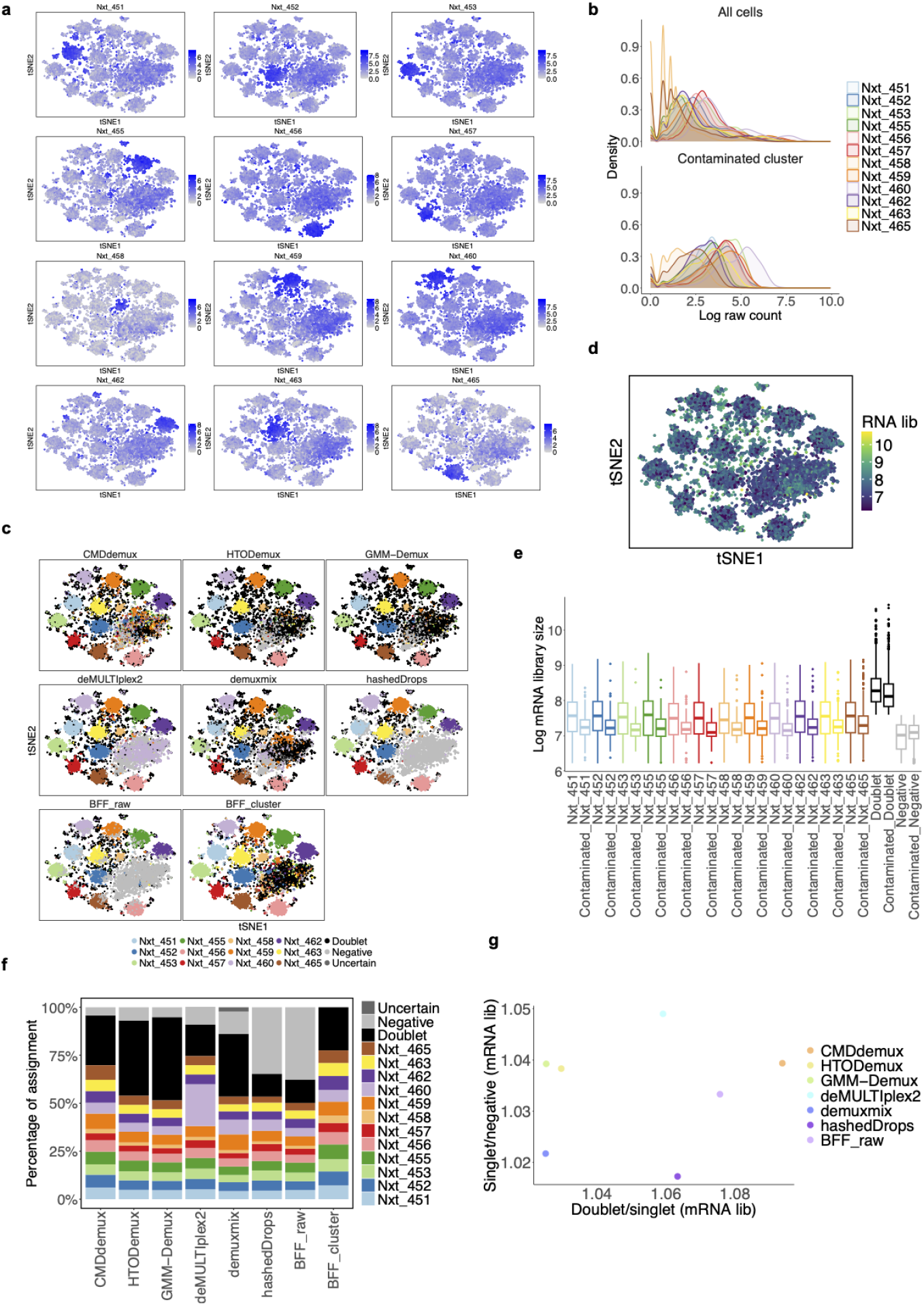
Performance of CMDdemux on the EMBRYO CMO dataset. (a) tSNE plot colored by the expression of each hashtag (log-transformed raw counts). (b) Distribution of log-transformed counts of each hashtag across all cells (top panel) and across cells in the contaminated cluster (bottom panel). (c) tSNE plot colored by demultiplexing results from different methods. demuxEM fails to output demultiplexing results. (d) tSNE plot colored by mRNA library sizes. (e) Comparison of mRNA library sizes between cells in the contaminated cluster and the corresponding demultiplexing categories in non-contaminated clusters. (f) Proportion of cells assigned to different demultiplexing categories by each method. (g) The x-axis shows the ratio of median mRNA library sizes between doublets and singlets (specifically, the singlet category with the largest median mRNA library size). The y-axis shows the ratio of median mRNA library sizes between singlets (the category with the smallest median mRNA library sizes) and negatives.

Cross-contamination can be caused by multiple factors, and one possible reason is exchange of hashtags or lipid oligos after sample pooling. The time after sample washing and pooling but before loading on the 10x Genomics instrument can be several minutes, in this period signal becomes contaminated in a time dependent fashion[25]. Another possible reason is inadequate washing. Although sufficient washing is required by the protocol, excessive washing may lead to cell loss. To preserve more cells, some experiments may be conducted with fewer washes, which can result in residual unbound antibodies. These free-floating antibodies may non-specifically bind to cells, leading to contaminated labelling. In addition, the MULTI-seq demultiplexing method is generally less stable than antibody-based hashing techniques, as it places higher demands on cell quality—including cell viability and debris—and is more prone to issues like fluid lipid contamination[28].

### CMDdemux enables handling of data with clusters of empty droplets

Empty droplets are common in single-cell data. These are droplets that do not contain a cell but still capture some ambient RNA transcripts[29]. Since these droplets lack cells, they are not labelled by hashtags. When present in large numbers, empty droplets can form a distinct cluster of negative cells. Generally, the hashtag quality in this data type is high. The main difference between this data and other high-quality datasets is that negative cells are fewer in number and tend to be scattered throughout the singlet cluster in the high-quality data. During cell clustering, an additional step is required to isolate the cluster of empty droplets, as described in Supplementary Methods Section S2. The empty droplet cluster is also distinct from the contaminated cluster: cells in the empty cluster are not labelled with any hashtags and exhibit low HTO and mRNA counts, whereas cells in the contaminated cluster show high HTO counts due to multiple hashtag attachments and include a mixture of negatives, singlets, and doublets.

In the PBMC data[1], there is an extra cluster (Cluster 1) (Fig. 9a) characterized by both low HTO library sizes and low mRNA library sizes (Fig. 9b), which corresponds to empty droplets. Each hashtag is highly expressed in a major singlet cluster, except in the empty cluster, which does not show expression of any hashtag (Fig. S8a) and exhibits low counts across all hashtags (Fig. S8b). All methods show high concordance in demultiplexing (Fig. 9c), except for BFF cluster, which incorrectly assigns the empty cluster as doublets, contradicting the truth. Compared to other methods, CMDdemux produces similar proportions of cells from different samples (Fig. S8c) and yields a reasonable doublet and negative rate (Fig. S8d). CMDdemux shows minor disagreements in assigning subclusters around the major singlet clusters. Specifically, CMDdemux classifies some cells in these subclusters as singlets, whereas other methods assign them as doublets. To further investigate, we examined these additional singlets defined by CMDdemux. Although these singlets exhibit higher HTO library sizes—similar to doublets (Fig. S8e)—they show mRNA library sizes comparable to their respective singlet categories and substantially lower than those of doublets (Fig. 9d). These findings support the advantage of CMDdemux, which leverages both cell hashing and gene expression data, offering more accurate demultiplexing than methods relying on cell hashing alone. Using several clustering-based metrics, including the silhouette score, Davies–Bouldin (DB) index, Dunn index, and Calinski–Harabasz (CH) index, CMDdemux demonstrates performance comparable to other methods (Fig. S8f, S8g).

**Fig. 9.**
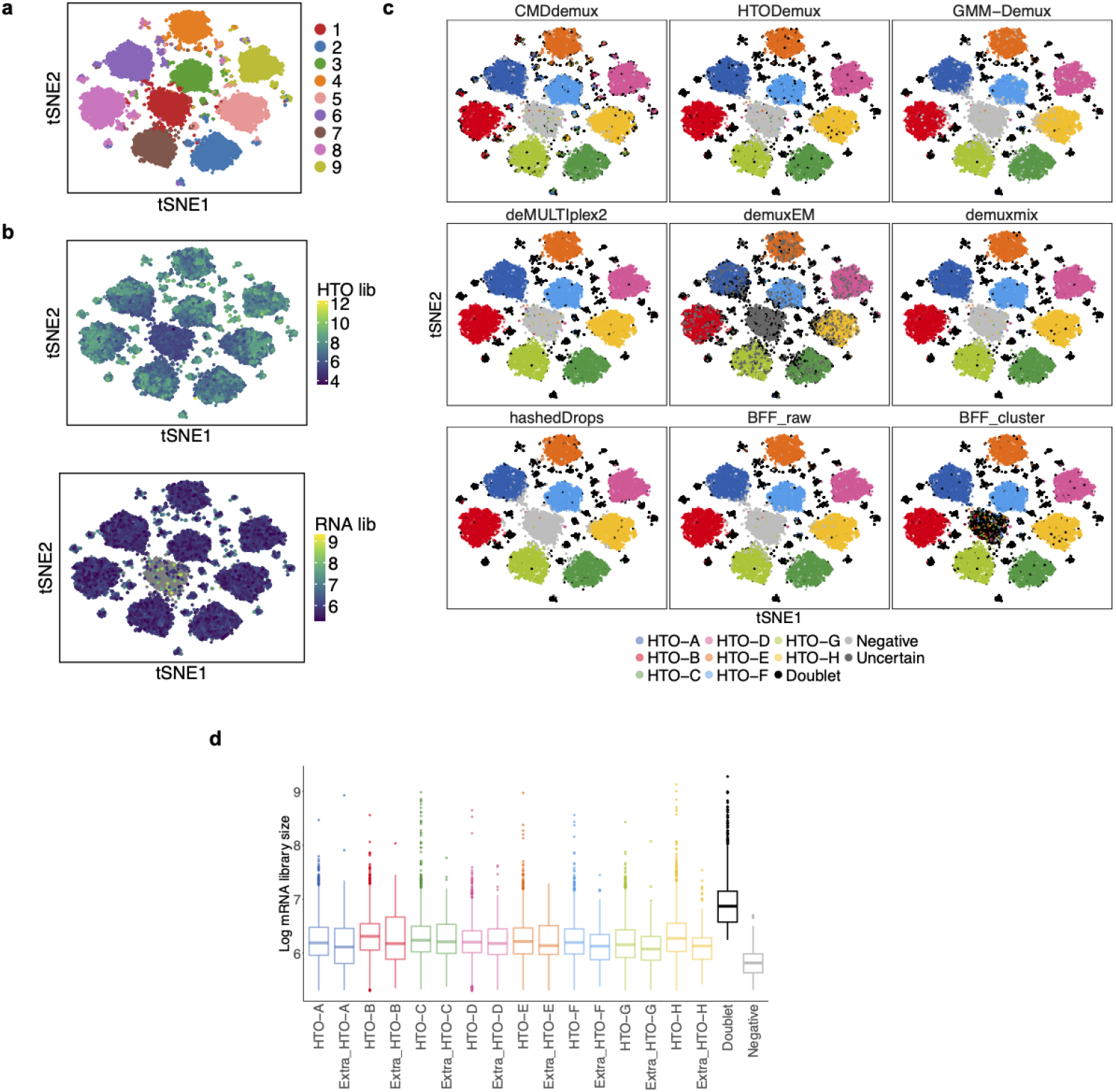
Performance of CMDdemux on PBMC data. (a) t-SNE plot colored by clustering of cell hashing data. (b) t-SNE plot colored by HTO library sizes (top panel) and mRNA library sizes (bottom panel). Grey dots represent low-quality cells that have been filtered out. (c) t-SNE plot colored by demultiplexing results from different methods. (d) Log-transformed mRNA library sizes of cells in different demultiplexing categories defined by CMDdemux. Extra singlets refer to cells identified as singlets by CMDdemux but as doublets by HTODemux.

It is apparent that the formation of the empty droplet cluster is related to cell quality. These cells are likely dead or have low viability, leading to disrupted cell membranes that prevent proper labelling by hashtags. Another possible reason is related to droplet capture inefficiency—cells may be labelled but not successfully encapsulated in droplets due to the stochastic nature of droplet formation. Since these cells are not captured within droplets, the absence of cell hashing signals is not surprising.

### CMDdemux improves performance on low-input data

Low-input data is also a common type of low-quality data, often resulting from failed cell hashing experiments. In general, low-input datasets exhibit low cell hashing counts across the entire cell population. Since all cells have low hashing counts, it is difficult to obtain clearly separated clusters corresponding to different hashtags. Instead, cells tend to clump together without distinct cluster boundaries. Due to the low hashing signal, defining thresholds to classify cells as negatives, singlets, or doublets becomes challenging. Low-input data differs from data with weak signals: in low-input data, all hashtags show weak signals across all cells, whereas weak-signal datasets typically involve only a few unevenly labelled hashtags affecting a subset of cells.

The PDX Hashtag Ab dataset[25] is a typical example of low-input data. All hashtags show low expression levels across all cells (Fig. S9a) and exhibit similar count distributions with weak signals (Fig. 10a). Moreover, the distributions of each hashtag are not clearly bimodal (Fig. S9b), making it difficult to distinguish signal from noise. Cells exhibit not only low HTO library sizes but also low mRNA library sizes (Fig. 10b). BFF and demuxEM fail to work on this low-input dataset, while deMUL-TIplex2 and hashedDrops produce suboptimal results, with a large proportion of cells classified as negatives (Fig. 10c). CMDdemux, HTODemux, and GMM-Demux assign a more appropriate number of singlets compared to the other methods (Fig. 10d), with CMDdemux achieving the lowest doublet and negative rates among the three (Fig. S9c).

**Fig. 10.**
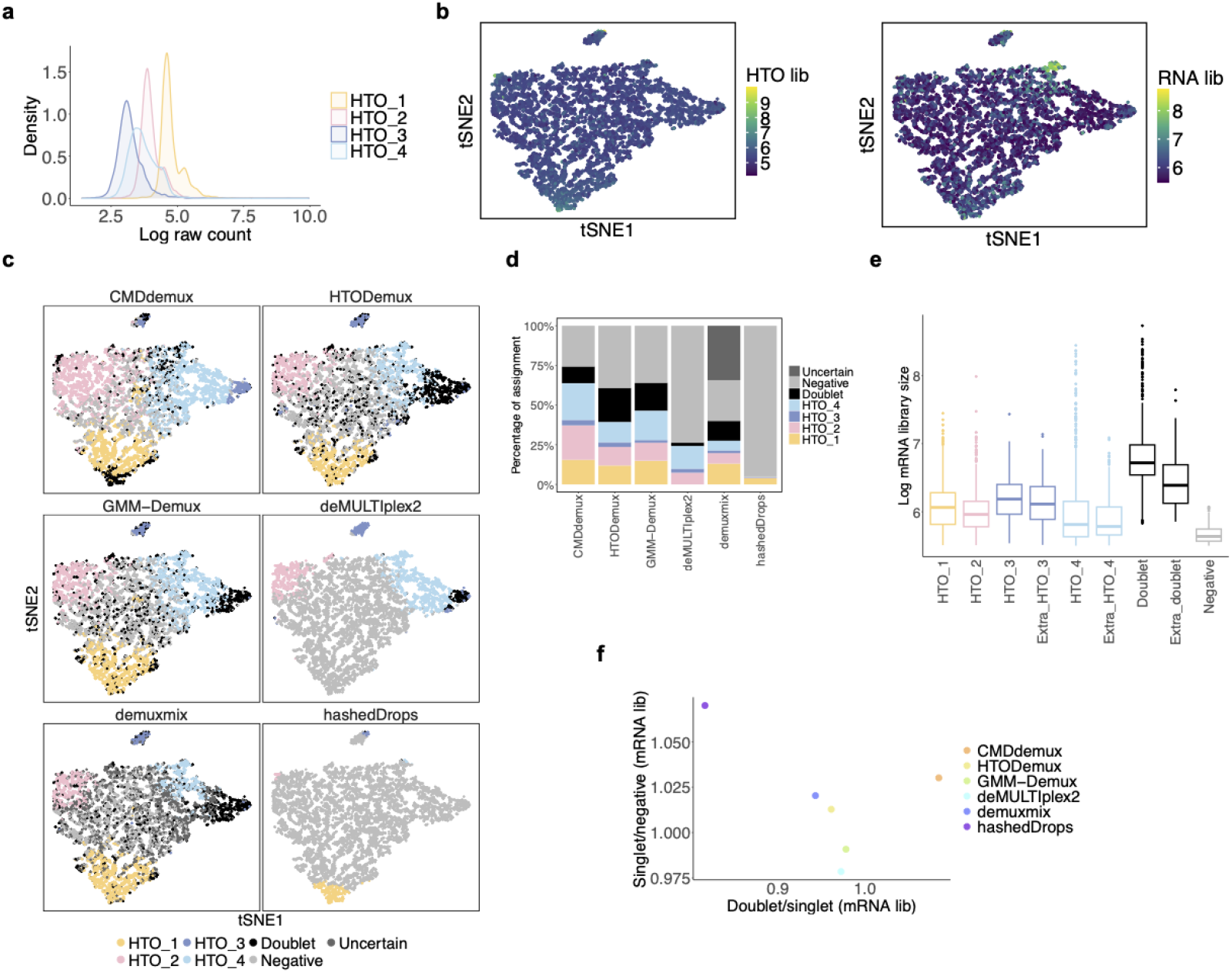
Performance of CMDdemux on PDX Hashtag Ab data. (a) Distribution of raw hashtag counts on a log scale. (b) t-SNE plots colored by HTO library sizes (left panel) and mRNA library sizes (right panel). (c) t-SNE plots colored by demultiplexing results from different methods. demuxEM and BFF are unable to generate demultiplexing results. (d) Proportion of cells assigned to different demultiplexing categories by each method. (e) mRNA library sizes of cell categories assigned by CMDdemux. “Extra HTO 3” and “Extra HTO 4” refer to cells assigned as HTO 3 and HTO 4 singlets by CMDdemux but as doublets by HTODemux and GMM-Demux. “Extra doublet” refers to cells assigned as doublets by CMDdemux but as HTO 1 singlets by both HTODemux and GMM-Demux. (f) Ratio of median mRNA library sizes between doublets and singlets (specifically, the singlet category with the largest median mRNA library size), as well as between singlets (with the smallest median mRNA library size) and negatives.

The major disagreements between CMDdemux and the other two methods occur in droplets within the HTO 1 cluster. CMDdemux classifies these droplets as doublets, whereas the other two methods classify them as HTO 1 singlets. Additionally, some cells in the HTO 4 cluster are assigned as HTO 3 or HTO 4 singlets by CMDdemux, but as doublets by the other methods. The additional doublets assigned by CMDdemux tend to have higher mRNA library sizes, while the additional singlets tend to have lower mRNA library sizes (Fig. 10e). Furthermore, these extra cells assigned by CMDdemux are correctly located within the doublet and singlet regions, respectively (Fig. S9d). Overall, CMDdemux-demultiplexed cells show a clear separation of mRNA library sizes among singlets, doublets, and negatives (Fig. S9e), and CMDdemux exhibits a high ratio of median mRNA library size between doublets and singlets, as well as between singlets and negatives (Fig. 10f).

The PDX MULTI-Seq CMO data[25] is another example of low-input data, characterized by overall low expression levels across all hashtags (Fig. 11a, b). The mRNA library sizes of these cells are also low (Fig. 11b). Additionally, this dataset exhibits uneven labelling, with stronger signals for Nxt 451 and Nxt 452. This is evident from the right-shifted distributions of their hashtag counts compared to the other two hashtags (Fig. 11a), and their higher expression levels across all cells (Fig. S10a). The unimodal distributions of these two hashtags further complicate demultiplexing (Fig. S10b). GMM-Demux fails to demultiplex Nxt 451 and Nxt 452 singlets (Fig. 11c), likely because the distributions of these hashtags violate the assumption of bimodality. deMULTIplex2, hashedDrops, and BFF raw fail to demultiplex three out of four hashtags. In contrast, CMDdemux, HTODemux, and demuxmix successfully demultiplex all four hashtags, although the latter two classify fewer Nxt 451 and Nxt 452 singlets compared to CMDdemux (Fig. 11c, d). Upon further examination of the additional Nxt 451 and Nxt 452 singlets assigned by CMDdemux, we observe that their mRNA library sizes are comparable to those of other singlet categories and are larger than those of negatives (Fig. 11e). Moreover, these extra singlets are located in the singlet region (Fig. S10c).

**Fig. 11.**
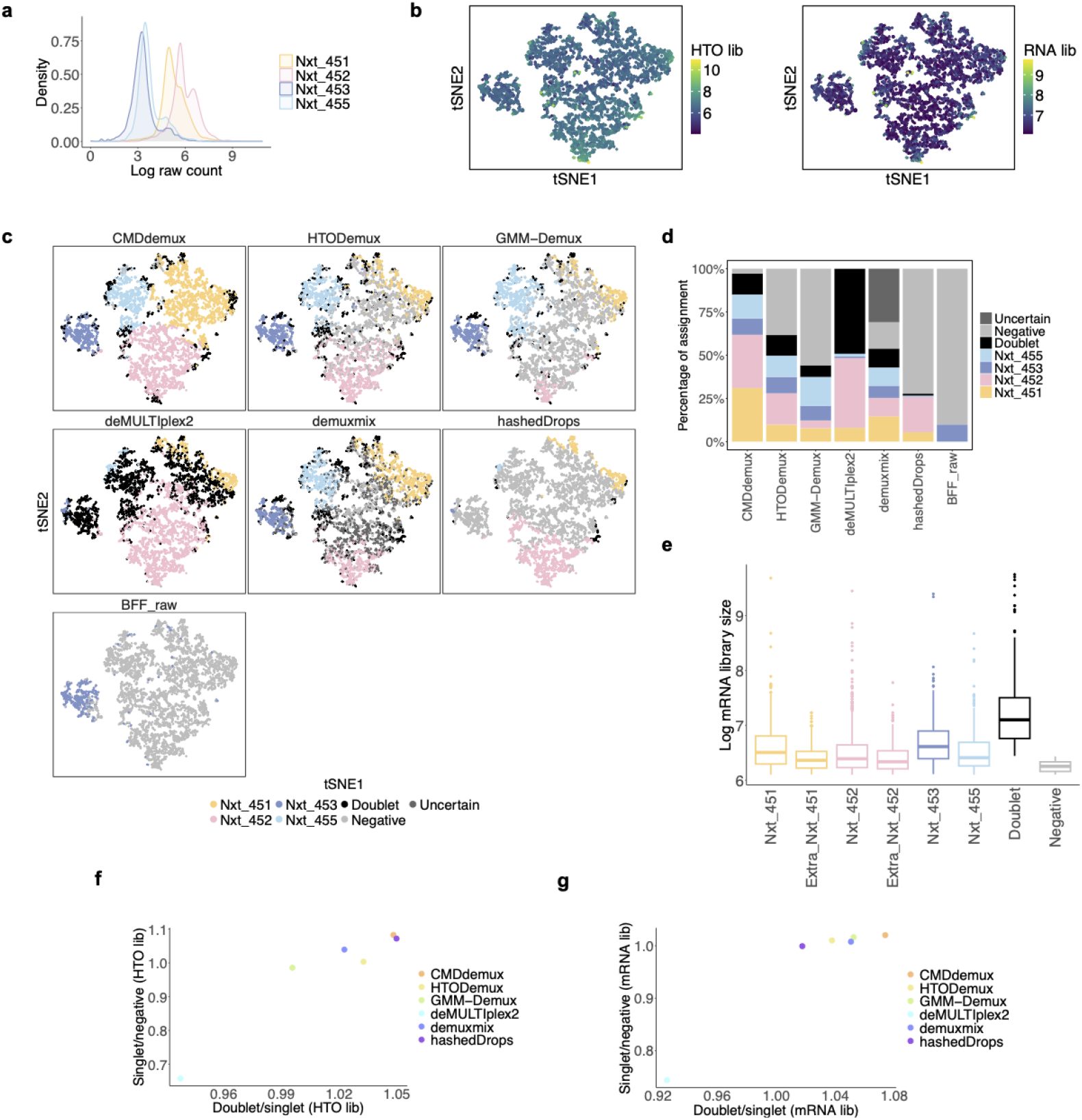
Performance of CMDdemux on PDX MULTI-Seq CMO data. (a) Distribution of raw hashtag counts on a log scale. (b) t-SNE plots colored by HTO library sizes (left) and mRNA library sizes (right). (c) t-SNE plots colored by demultiplexing results from different methods. demuxEM and BFF cluster are unable to produce demultiplexing results. (d) Percentage of cells assigned to different demultiplexing categories across methods. (e) Log-transformed mRNA library sizes of different demultiplexing categories assigned by CMDdemux. The “Extra Nxt 451” and “Extra Nxt 452” categories refer to additional Nxt 451 and Nxt 452 singlets assigned by CMDdemux but classified as negatives by both HTODemux and demuxmix. (f) Ratio of median HTO library sizes between doublets and singlets (the singlet category with the highest median HTO library size), and between singlets (with the lowest HTO library size) and negatives. (g) Ratio of median mRNA library sizes between doublets and singlets (the singlet category with the highest median mRNA library size), and between singlets (with the lowest mRNA library size) and negatives.

Overall, CMDdemux rescues the most singlets compared to all other methods (Fig. 11d), preserving the highest number of cells for downstream analysis. CMDdemux also achieves a reasonable doublet rate and a low negative rate (Fig. S10d). The mRNA library sizes assigned by CMDdemux are clearly separated among singlets, doublets, and negatives (Fig. S10e). Moreover, CMDdemux-demultiplexed cells exhibit both high median HTO library size ratios and high median mRNA library size ratios—between doublets and singlets, as well as between singlets and negatives (Fig. 11f,g; Fig. S10f,g)—further supporting CMDdemux as a robust method for demultiplexing low-input data.

The major cause of low-input data is the technical difficulty of single-nucleus experiments. This dataset originates from such experiments, where nuclei were extracted and labelled. Single-nucleus protocols are often used when samples are difficult to freeze or dissociate[30]. However, compared to single-cell hashing, labelling nuclei is more challenging. This is because the cytoplasm is disrupted during extraction, allowing antibodies to non-specifically bind to cytoplasmic debris and lysed cellular fragments[3]. In the case of the PDX data, tumour nuclei are inherently fragile. Inadequate incubation and washing steps can result in reduced antibody binding.

Conversely, more aggressive processing steps to improve labelling may cause cross-contamination or loss of nuclei, further lowering data quality. Additionally, according to the original publication[25], the nuclei in the PDX samples were in good condition immediately after isolation, but their quality deteriorated following the hashing process—contributing to the low-input nature of the data.

### CMDdemux provides reliable demultiplexing results comparable to other methods on high-quality data

All methods perform well on high-quality data, as their models are designed based on such data. This study includes three datasets—human brain[3], BAL[31], and vehicle mouse[24]. It is evident that each hashtag is uniquely and highly expressed in one cluster of high-quality cells (Fig. S11a, S12a, S13a), and the expression of each hashtag follows a standard bimodal distribution (Fig. S11b, S12b, S13b). The distributions of hashtags are also similar across samples (Fig. S11c, S12c, S13c). The HTO and mRNA library sizes of the high-quality data fall within a reasonable range (Fig. S11d, S12d, S13d). For high-quality data, all methods yield very similar demultiplexing results (Fig. 12a,d,g). The proportion of cells assigned to each donor by CMDdemux closely matches those from other methods (Fig. S11e, S12e, S13e), especially for the human brain and BAL datasets, where CMDdemux results align well with SNP-based demultiplexing. CMDdemux also outputs an approximately equal proportion of singlets from different donors in these two datasets.

**Fig. 12.**
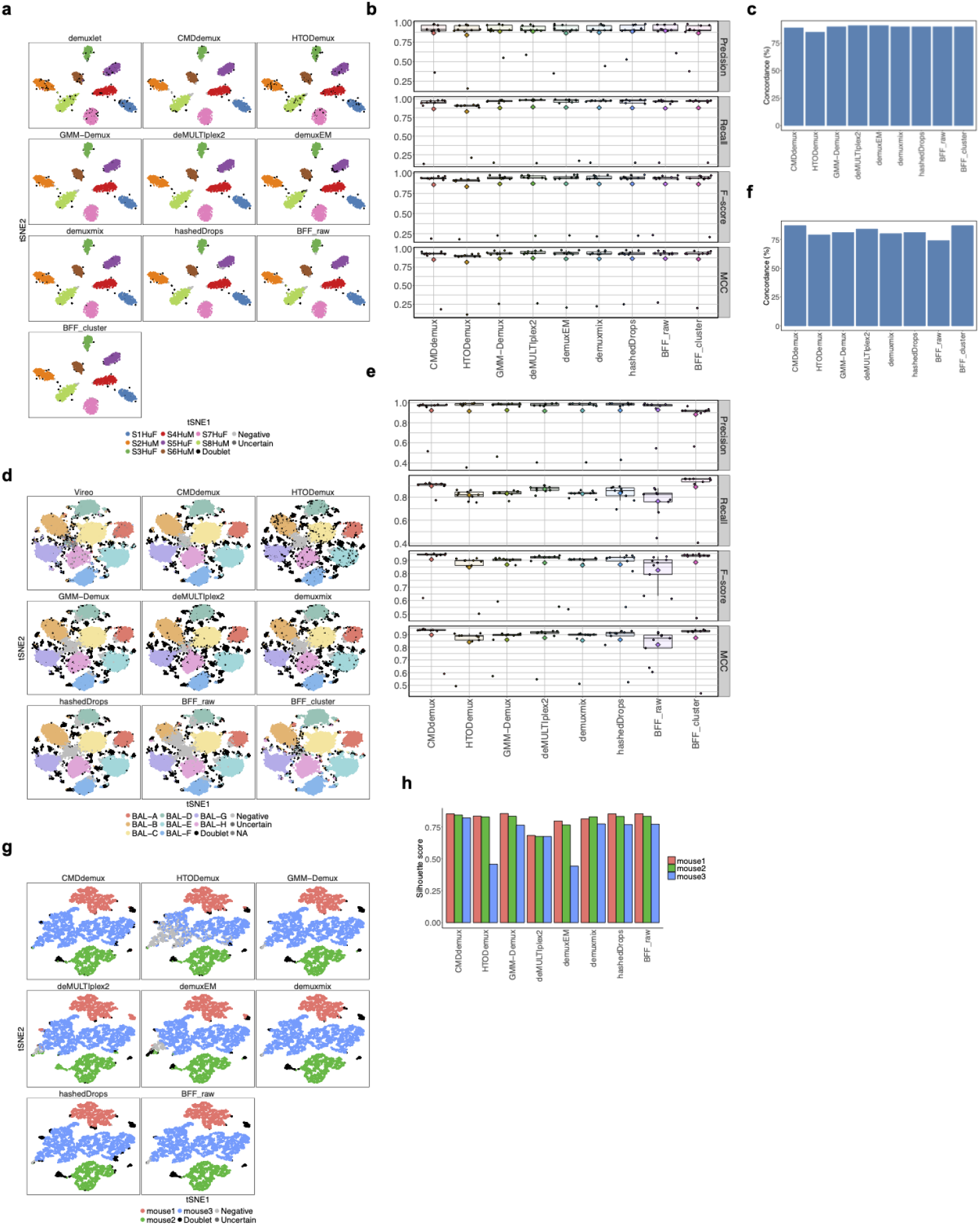
Performance of CMDdemux on high-quality data. (a–c) Demultiplexing results for human brain data. (a) t-SNE plot colored by demultiplexing results from different methods. (b) Performance evaluation using precision, recall, F-score, and MCC, with demuxlet used as the ground truth. (c) Overall concordance of different methods with demuxlet. (d–f) Demultiplexing results for BAL data (gene expression data not available). (d) t-SNE plot colored by demultiplexing results from different methods. For some cells, Vireo results are not available and are labelled as NA. demuxEM failed to generate demultiplexing results. (e) Performance evaluation using precision, recall, F-score, and MCC, with Vireo used as the ground truth. (f) Overall concordance of different methods with Vireo. (g, h) Demultiplexing results for vehicle mouse data. (g) t-SNE plot colored by demultiplexing results from different methods. BFF cluster failed to generate demultiplexing results. (h) Median silhouette score of each singlet category for different methods.

In these high-quality datasets, CMDdemux demonstrates reliable performance, as evidenced by its high concordance with SNP-based demultiplexing results both overall (Fig. 12c,f) and within each subcategory (Fig. S11f, S12f). CMDdemux also achieves high demultiplexing accuracy, as evaluated by precision, recall, F-score, and MCC (Fig. 12b,e; Fig. S11g; Fig. S12g). For the mouse vehicle dataset, where SNP-based demultiplexing is unavailable, CMDdemux achieves high silhouette scores across all three samples (Fig. 12h; Fig. S13f). Furthermore, CMDdemux-demultiplexed singlets, doublets, and negatives show clearly distinct mRNA library sizes, ranging from high to low, respectively (Fig. S11h; Fig. S13g,h,i).

### Local CLR normalization is more suitable than global CLR normalization for processing low-quality cell hashing data

CMDdemux uses a local CLR strategy, whereas HTODemux and GMM-Demux apply their methods based on global CLR normalization. The details of local CLR normalization are discussed in the Methods section, and global CLR normalization is described in Supplementary Methods Section S1. The major difference between the two normalization methods is that local CLR performs normalization within each cell, which can be interpreted as normalizing the hashing count of a given label relative to all other labels within the same cell. In contrast, global CLR normalization is applied across cells, meaning it normalizes the hashing count of a given cell relative to all other cells within the same hashing label. Another key difference is that global CLR directly takes the log ratio of the raw hashing count to the geometric mean of raw HTO counts across cells. In contrast, local CLR first performs library size normalization and then computes the log ratio of the normalized count to the geometric mean of that cell’s normalized counts across all HTOs. This extra normalization step in local CLR helps reduce the impact of library size differences.

There are two major consequences of these differences. First, global CLR tends to unify the distribution of hashtags, whereas local CLR preserves the differences among hashtags (Fig. 13a). The advantage of the unified distribution in global CLR is that it simplifies threshold determination, as all hashtags have similar distributions. While this strategy works well on high-quality data, it fails on low-quality data: in such cases, low-quality hashtags still differ slightly from high-quality ones even after global normalization and often do not form a bimodal distribution. Second, local CLR centers values around 0 (Fig. 13b–e). In local CLR-normalized space, cells form clusters, with major clusters representing cells from different donors, and subclusters or cluster edges representing doublets or negatives (Fig. 13b,c). In global CLR space, singlets from each donor tend to align along distinct axes (Fig. 13b,c), while doublets appear off-axis and negatives cluster near the origin (lower left corner). In the high-quality vehicle mouse data, both global and local CLR normalization meet the assumptions for proper demultiplexing (Fig. 13b). However, in the low-quality treated mouse data, only local CLR preserves the expected separation of singlets, doublets, and negatives. In contrast, cells are scattered across the global CLR space and no longer align with the expected axis directions (Fig. 13c).

**Fig. 13.**
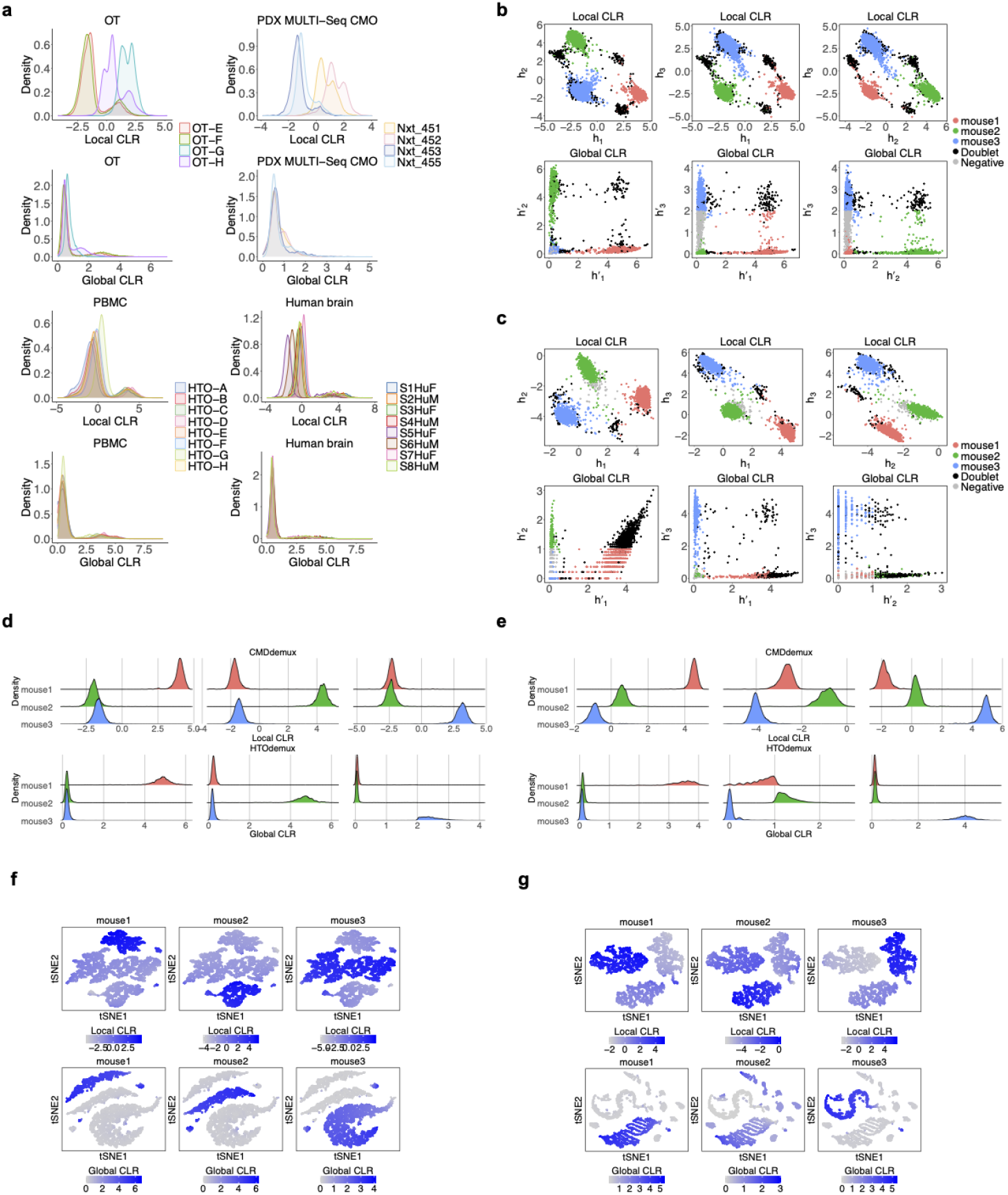
Comparison between local CLR and global CLR normalization. (a) Distributions of local CLR and global CLR values in OT (low-quality data), PDX MULTI-Seq CMO (low-quality data), PBMC (low-quality data), and human brain data (high-quality data). (b, c) Pairwise plots of local CLR-normalized values (h) or global CLR-normalized values (h’). Each dot represents a cell. In the local CLR space, cells are colored by CMDdemux results; in the global CLR space, cells are colored by HTODemux results. (b) Vehicle mouse data. (c) Treated mouse data. (d, e) Ridge plots showing the distribution of local CLR-normalized values (upper panels) and global CLR-normalized values (lower panels) for each singlet category. Each column corresponds to a singlet category demultiplexed by CMDdemux (upper panels) or HTODemux (lower panels). For example, the first panel in (d) shows the distribution of local CLR-normalized values for all mouse1 singlets identified by CMDdemux. (d) Vehicle mouse data. (e) Treated mouse data. (f, g) tSNE plots of each hashtag based on local CLR expression (upper panels) and global CLR expression (lower panels). The tSNE embeddings are computed from local CLR values (upper panels) and global CLR values (lower panels). (f) Vehicle mouse data. (g) Treated mouse data.

Both global CLR and local CLR work well on high-quality data, but their performance differs on low-quality data. In the high-quality vehicle mouse dataset, both methods clearly separate cells from the three mouse samples (Fig. 13d). However, in the low-quality treated mouse dataset, global CLR fails to distinguish mouse2 cells from mouse1 cells, as they exhibit similar global CLR expression levels (Fig. 13e). In contrast, local CLR provides more meaningful low-dimensional embeddings in lowquality data (Fig. 13f,g). In the high-quality dataset, both normalization methods produce three distinct clusters in tSNE space, corresponding to the three mouse samples, with each hashtag being uniquely and highly expressed in one cluster (Fig. 13f). In the low-quality treated dataset (Fig. 13g), global CLR fails to generate clearly separated clusters, and the mouse2 hashtag is moderately expressed across multiple clusters. In contrast, local CLR still yields three well-separated clusters, each with a uniquely and highly expressed hashtag.

## Discussion

Accurate demultiplexing plays an essential role in downstream analysis, as it determines the number of usable cells for subsequent steps. The sample-of-origin classification directly defines biological conditions for comparison—such as disease vs. healthy, drug-treated vs. control, or knockout vs. wild-type. Misclassified cells can lead to incorrect conclusions. CMDdemux significantly improves demultiplexing accuracy, especially for low-quality data. First, unlike other methods that offer a single all-in-one function, CMDdemux provides a clear, step-by-step workflow. While all-in-one functions may be convenient, they often fail with low-quality data, which may not fit the model assumptions or default parameters. CMDdemux allows users to examine each step carefully, helping to detect and avoid artifacts caused by low-quality inputs. Second, CMDdemux offers tools for inspecting suspicious droplets. In the absence of ground truth, users can visually assess questionable classifications using 2D and 3D plots, thereby improving confidence in the results. Third, CMDdemux enhances the accuracy of identifying negatives and doublets by incorporating information from mRNA profiles. Hashing-based demultiplexing alone provides limited insight, but CMDdemux reclassifies droplets using mRNA library sizes to produce more reliable results. Fourth, CMDdemux uses local CLR normalization, which preserves the unique characteristics of each hash and is more effective for low-quality data than global CLR, which tends to overly smooth the distribution and obscure true signals. Fifth, CMDdemux does not assume a bimodal distribution, making it robust to the irregular distributions found in low-quality data. The use of Mahalanobis distance provides a more flexible and accurate way to determine sample origin. CMDdemux also allows users to adjust the number of clusters and choose between medoids or median expression levels for sample assignment, adding flexibility when dealing with problematic datasets. Moreover, CMDdemux rescues more usable singlet cells. Competing methods often misclassify these as negatives or doublets. This is especially important when analyzing rare cell types, where preserving every informative cell is critical. Finally, CMDdemux assigns sample-of-origin information to doublets—something many methods overlook. This feature is particularly valuable for multi-hash experiments, where a single sample may be labelled with multiple hashtags. CMDdemux can distinguish and trace these back to their original donors.

CMDdemux performs well on both high-quality and low-quality data. In highquality data, where hashtag labelling is optimized and signals are strong, most demultiplexing methods perform well because the data fits their underlying assumptions. However, CMDdemux stands out by maintaining consistent and robust performance even in low-quality data. Many existing methods assume that hashtag signals follow similar distributions. In contrast, low-quality data often suffer from uneven labelling, leading to shifted distributions. For example, in datasets with weak signals, the background noise level of standard hashes may correspond to true signals in low-quality hashes. Conversely, in datasets with overly strong signals, the true signal of standard hashes may be misinterpreted as noise. Because most methods cannot distinguish between noise and true signals in these cases, they tend to misclassify such cells as negatives or doublets. CMDdemux, however, accurately demultiplexes these unevenly labelled cells as singlets. In contamination-prone datasets with tri-modal distributions, most models fail due to their bimodal distribution assumptions. Only CMDdemux and BFF_cluster_ correctly identify the contaminated cluster as a mixture of negatives, singlets, and doublets. In datasets containing empty droplet clusters (characterized by low HTO and mRNA library sizes), all methods perform similarly, but CMDdemux disagrees with others on a subset of doublets. By incorporating mRNA profiles, CMDdemux more accurately reclassifies these droplets as singlets. Low-input datasets show reduced expression across all hashtags. Methods like BFF, DemuxEM, and deMULTI-plex2 fail to perform in such scenarios. Other tools tend to over-call negatives, while CMDdemux recovers more singlets that can be retained for further analysis. Overall, CMDdemux generates a balanced demultiplexing outcome across donors, supporting the experimental assumption that donor cell counts should be similar at the start. DemuxEM performs inconsistently and often fails to produce results on low-quality data. hashedDrops frequently overcalls negatives, likely due to its conservative singlet threshold. deMULTIplex2 struggles with low-count hashing data, possibly because the GLM-NB model fails to fit sparse counts and the EM algorithm does not converge. BFF methods fail in some low-quality datasets, likely due to their reliance on bimodal distribution assumptions and rigid thresholding. HTODemux, GMM-Demux, and Demuxmix show only moderate performance, often misclassifying many cells from low-quality hashes as negatives.

Although CMDdemux outperforms other methods on low-quality data, it still has several limitations. First, some manual thresholds must be set by users. While we provide default parameters for each step and automatic settings generally work well in most datasets, we still recommend that users examine the data carefully. CMDdemux includes built-in functions to help visualize the data and guide parameter selection. One key reason for the failure of other methods is their all-in-one structure, which makes it difficult to troubleshoot errors in low-quality data. Because the distribution of hash counts can vary across different low-quality scenarios, inspecting the data and adjusting parameters is essential for achieving optimal results. Second, it can sometimes be challenging to clearly distinguish outlier cells, negatives, and doublets from singlets when their Mahalanobis distances are close to the classification cutoff. Since these groups may have similar distances, Mahalanobis distance alone may not perfectly separate them. If users are uncertain about the demultiplexing results, CMDdemux provides 2D and 3D visualization tools to inspect droplets near the decision boundary. Third, CMDdemux cannot identify doublets formed by cells from the same sample or negative droplets containing genuine cells that lack hash labelling. These are limitations of the multiplexing technology itself rather than limitations of CMDdemux, as such droplets present hashing profiles indistinguishable from singlets and negatives, respectively. All demultiplexing algorithms, not just CMDdemux, will classify these droplets as singlets or negatives. However, CMDdemux improves upon other methods by using mRNA profiles to reclassify negatives and doublets, providing more accurate classifications than methods relying solely on hashing data. Finally, in rare cases involving unlabelled hashes, the thresholds for log expression and CLR differences are empirically set. This condition is uncommon; in fact, EMBRYO LMO is the only dataset encountered with this issue. It likely indicates a failure in the labelling process for the affected hash. In such cases, users are encouraged to tune parameters carefully if they wish to recover cells associated with the unlabelled hash.

The quality of hashing data is highly dependent on the experiment. Although each instance of low-quality data may have its own specific causes, there are some common reasons for poor-quality results. First, cell state is crucial in determining the binding efficiency of hashtags. For example, drug treatment may alter cell states and affect their binding to hashtags. Cancer cells can be difficult to label at different progression stages compared to healthy cells. Nuclei are more fragile and prone to degradation compared to other samples, requiring extra careful processing. Furthermore, the choice of protocol can also impact data quality. Antibody-based techniques generally perform more reliably than lipid-based labelling strategies[16]. However, it should be noted that antibodies may show cell-type preferences, especially for immune cells. In addition, the procedures during the hashing experiment are critical. For example, the washing step is frequently emphasized in hashing workflows because residual hashes in the buffer can worsen signal quality. Incubation time also influences binding efficiency. Taken together, improving experimental design and hashing procedures is essential to enhance the quality of hashing data.

In summary, CMDdemux demultiplexes droplets using the Mahalanobis distance in a space after within-cell CLR normalization. CMDdemux exhibits superior performance on both high- and low-quality data. It is capable of demultiplexing cells generated from various technologies and handling diverse types of low-quality hashing data. CMDdemux helps rescue singlets from donors that are poorly labelled without compromising accuracy, which may retain more cells from rare cell types or delicate samples, leading to more meaningful downstream analysis.

## Methods

### Local CLR normalization

In order to distinguish the across-cell CLR normalization method from HTODemux and GMM-Demux, our within-cell CLR normalization is named as local CLR normalization. To perform local CLR, the HTO count for hashtag *i* and cell *c* is initially normalized by the cell’s HTO library size. The library size normalized HTO counts *π*_*i,c*_ for *N* hashtags are

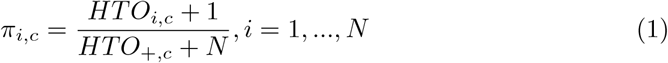

where *HTO*_+,*c*_ = Σ_*i*_ *HTO*_*i,c*_. To deal with zero counts, we add a small pseudo value 1 to each entry of the HTO count matrix. Then the geometric mean across hashtags for each cell is calculated as:

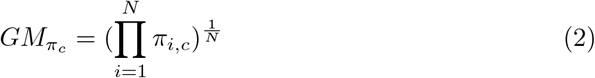

The log of the geometric mean scaled library size normalized HTO value is the local CLR value *h*_*i,c*_.

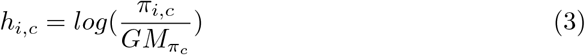

From now on, local CLR normalized HTO data will mean this quantity.

### K-medoids clustering

Unsupervised K-medoids clustering aims to group cells labelled by the same hashtag. The default set is *K* = *N* so that theoretically each cluster can represent a hashtag. The centroid *M* ^(*l*)^ of each cluster *l* is also available in this step. Considering the speed of the clustering algorithm, we provide two modes: the pam function[32] is recommended for datasets with fewer than 10,000 cells; for larger datasets, the faster clara function[33] is recommended.

For high-quality data, setting the number of clusters equal to the number of hashtags typically performs well. For low-quality data, however, the choice of cluster number can significantly affect demultiplexing accuracy. To assist users in selecting the appropriate number of clusters, we provide a visual inspection method: box plots of local CLR expression for each hashtag across clusters. An ideal value of *k* is one in which each hashtag has a single cluster showing substantially higher local CLR values than the others. In addition to visual inspection, we offer a quantitative approach to determine the optimal number of clusters. For each hashtag, we compute the median local CLR value across all clusters and identify the top cluster (i.e., the one with the highest median CLR), defined as follows:

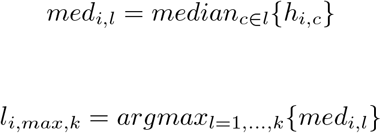

To compare two candidate cluster numbers, *k*_1_ and *k*_2_ (*k*_2_ *> k*_1_), we calculate the difference in median CLR values for each hashtag between their respective top clusters:

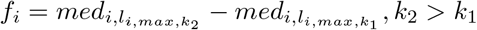

This difference can be interpreted as the log fold-change of hashtag expression between the top clusters for the two *k* values. If the maximum of *f*_*i,l*_ across all hashtags, i.e.,*max*_*i*,…,*N*_ {*f*_*i*_}, exceeds a given threshold (default: 1), then *k*_2_ is recommended for downstream analysis. An example is shown in Supplementary Methods Section S2.

### Defining core and non-core cells

Core cells are the cells closer to their cluster centroid, while non-core cells are farther from their cluster centroid. Core and non-core cells are defined by their Euclidean distances to the centroid of their cluster. The Euclidean distances for all cells within a cluster to its cluster centroid can be obtained, and a threshold set up using a quantile of the distance distribution. The default threshold is 0.9, which means that 90% of cells with smallest Euclidean distance to the cluster centroid are core cells, and the remaining 10% of cells with larger Euclidean distance are non-core cells. Sometimes this default threshold is not good enough to distinguish the core and non-core cells. Therefore, we recommend visualizing the distribution of the Euclidean distance and manually modifying this parameter if the default threshold does not work. For data containing clusters of negatives, all cells within those negative clusters will be classified as non-core cells. The main reason to define core and non-core cells is that non-core cells include background noise and multiplets and are outliers. In the following step, the matrix of covariances between each pair of HTOs is required, and we do not want outliers to influence the covariance matrix estimation.

### Labelling clusters by their sample of origins

The k-medoids clusters can be labelled either by the cluster medoids or by the average expression level. In the medoid-based approach, the local CLR values of each hashtag across cluster medoids are compared. A hashtag *i* is assigned to the cluster *l* whose medoid has the highest local CLR value, i.e., 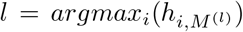. In the average expression-based approach, non-core cells are first excluded. The median local CLR value of each hashtag across cells in each cluster is then computed to represent the hashtag’s average expression. Hashtag *i* is assigned to the cluster *l* with the highest median CLR value: *l* = *argmax*_*i*_(*med*_*i,l*_). In general, the medoid-based method works well in most scenarios and is therefore set as the default. However, the average expression-based method is better suited for cases where conflicting hashtags are assigned to the same cluster. This scenario is further discussed in Supplementary Methods Section S3.

### Calculating Mahalanobis distances

Since non-core cells include outliers that influence the covariance matrix, we only include core cells to calculate the matrix of covariances Γ between pairs of local CLR normalized HTO values. Its entries are

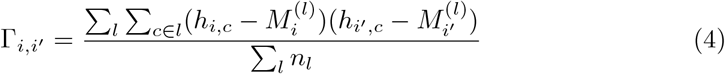

where *n*_*l*_ is the number of cells in cluster *l*.

The next step requires the inverse Γ^−1^ of the matrix Γ. However, Γ is singular and so not invertible. The following shows its singularity:

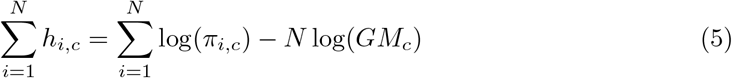

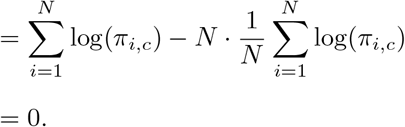

Equation 5 proves the linear dependence between log local CLR normalized HTO values. Since the columns of Γ are linearly dependent, the determinant of Γ is 0. Therefore, Γ is singular. We use the Moore-Penrose generalized inverse covariance matrix Γ^+^ to get the pseudo inverse of Γ.

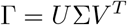

Σ is the diagonal matrix of non-zero singular values. *U* is the orthogonal matrix of left singular vectors, and *V* is the orthogonal matrix of right singular vectors.

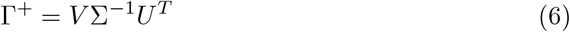

Then the Mahalanobis distance *MD*_*c,l*_ for each cell *c* to the centroid *M* ^(*l*)^ of the K-medoids cluster *l* is defined by:

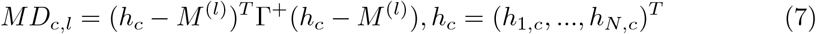

### Initial demultiplexing using Mahalanobis distances

An initial classification step assigns cells into singlets and outliers based on the distribution of minimum Mahalanobis distances across cells. Theoretically, the distribution of minimum Mahalanobis distances should be bimodal, with a large mode of smaller mean representing singlets (which have smaller Mahalanobis distances) and a smaller mode of larger mean representing negatives and doublets (which are more distant from the cluster centroid, reflected by larger Mahalanobis distances). However, this theoretical distribution does not always fit low-quality data. We recommend users set the cut-off based on a visual inspection of the distribution’s quantile. Two types of default cut-offs are provided. One is the 0.95 quantile of the observed distribution. The other is empirically based, assuming that squared Mahalanobis distances follow a chi-square distribution on *N* − 1 degrees of freedom, and sets the cut-off at the 0.975 quantile. Cells with minimum Mahalanobis distances smaller than the cut-off are classified as singlets, while others are classified as outliers. Singlet cells are directly assigned to the hashtag corresponding to their minimum Mahalanobis distance.

### Further distinguishing negatives and doublets among the outliers

We assume that the log HTO library size for outlier cells follows a bimodal distribution. This is because the outlier population includes both negative cells and doublets. Negative cells have too few HTO UMIs to reliably identify their samples, while doublets encapsulate multiple cells and therefore contain relatively more HTO UMIs within the gel beads. As a consequence, we expect one peak with smaller log HTO library sizes corresponding to negatives, and another peak with larger log HTO library sizes corresponding to doublets. The midpoint (antimode) between these two peaks is used as the threshold to distinguish negatives from doublets, which is determined using the locmodes function from the multimode package[34]. Additionally, the possible sample identities for doublets are inferred based on the hashtags with the smallest and second-smallest Mahalanobis distances. This information can assist in demultiplexing experiments where samples are labelled with mixed HTOs.

Since using only one modality for classification is not always reliable, we also utilize transcriptome data to reclassify outlier cells when gene expression data is available. Specifically, the median log mRNA library size across all singlets is used as a threshold.

Droplets initially classified as negatives but with mRNA library sizes larger than the threshold are reclassified as singlets. Similarly, droplets initially classified as doublets but with mRNA library sizes smaller than the threshold are also reclassified as singlets. This reclassification strategy helps prevent misclassification when relying solely on cell hashing data.

### Benchmarking on real datasets

The raw hashing counts from all datasets were used for benchmarking. Since CMDdemux requires some manual settings, all other methods were run using default settings. In the PDX CellPlex dataset, six cells could not be demultiplexed by deMULTIplex2 and were labelled as “Uncertain”. For demuxmix, datasets without gene expression information were analyzed using the “naive” mode. For hashedDrops, all droplets without a confident assignment were classified as “Negative”. For demuxEM, droplets marked as “unknown” were reclassified as “Negative”. The mRNA library size for each cell is defined as the sum of all counts from the scRNA-seq gene expression matrix *Y* : 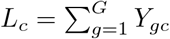where *g* denotes gene *g* in the rows of matrix *Y*.

To evaluate the performance of different methods, we computed the median mRNA or HTO library size ratios: between doublets and singlets; between singlets and negatives.

Specifically, we calculated the median mRNA library size 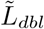 and HTO library size 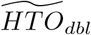 across all doublets. For singlets, we calculated the median mRNA library size 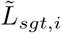 or HTO library size 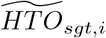 within each donor i, and identified the donor with the maximum median value 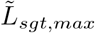 or 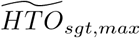. The doublet-to-singlet ratio was computed as:

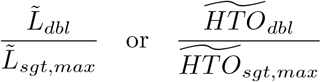

Similarly, for singlet-to-negative comparisons, we used the minimum median library size among donors:

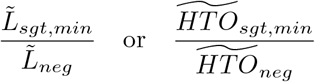

### Evaluation metrics on data without ground truth

The silhouette score was used to evaluate the separation of clusters[35]. The approxSilhouette function[36] was used to compute the silhouette score for each single cell. To avoid bias from negatives and doublets, only the silhouette scores of singlets were used for evaluation. A high silhouette score indicates good performance by a method, meaning that demultiplexed singlets form clearly separated clusters by donor. Another metric used was the Calinski–Harabasz (CH) index[37], which also measures clustering quality. The `index.G1` function[38] was used to calculate the CH index based on the log-transformed raw hashing counts using Euclidean distance. All doublets and negatives were excluded from the CH index calculation. A higher CH index indicates better performance of the demultiplexing method. The Davies–Bouldin (DB) index was calculated using the `index.DB` function[38], where a lower value corresponds to better clustering. The Dunn index was also calculated using the `dunn` function[39], with higher values indicating better performance.

### Evaluation metrics on data with ground truth

Demultiplexing results from SNP-based methods were used as ground truth to benchmark cell hashing-based methods. Results from Vireo and demuxlet were obtained from their respective original publications. Concordance between cell hashing-based and SNP-based methods was evaluated by calculating the number of cells assigned the same label in both methods, divided by the total number of cells. The overall concordance was computed as the sum of all category-level concordance values. In addition, we used precision (*P*), recall (*R*), F-score (*F*_1_), and Matthews correlation coefficient (*MCC*)[40] to evaluate the performance of each method. Define

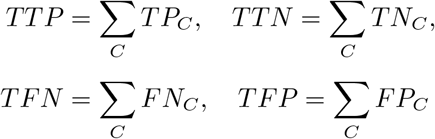

Then the micro-averaged values of these metrics were computed as follows:

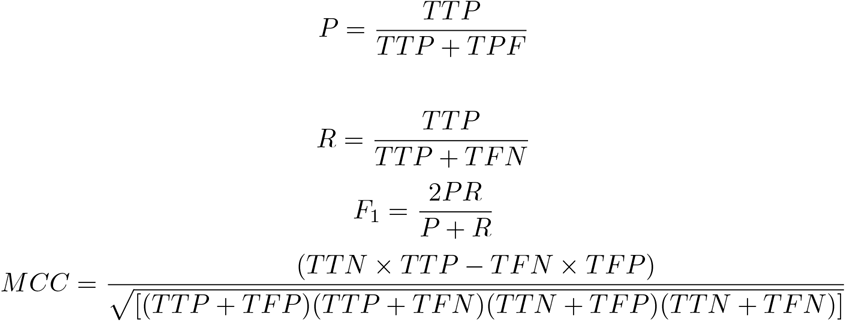

Here,

*C* denotes each demultiplexing category.

*TP*_*C*_ (true positives): cells classified into category *C* by both the ground truth and the benchmarking method.

*TN*_*C*_ (true negatives): cells not classified as *C* by either method.

*FP*_*C*_ (false positives): cells classified as *C* by the benchmarking method but not by the ground truth.

*FN*_*C*_ (false negatives): cells classified as *C* by the ground truth but not by the benchmarking method.

### Data

A summary of all benchmarking data is provided in Table S3.

#### Human brain

The human brain data[3] consists of eight dorsolateral prefrontal cortex samples, including four from females and four from males. The samples were labeled using anti-nuclear pore complex antibodies.

#### BAL

The BAL data[31] was derived from eight paediatric bronchoalveolar lavage (BAL) fluid samples labeled with TotalSeq-A antibodies. Only batch 1 samples were used. Gene expression data is not available for this dataset.

#### Mouse

The mouse data[24] includes samples from vehicle-treated and ipatasertib-treated mice. AT3OVA tumors were harvested after eight days of treatment with either vehicle or ipatasertib in three mice per group. Each mouse was labeled using TotalSeq™ anti-mouse hashtag antibodies. CD45^+^ cells were sorted using a BD FACS Fusion sorter. Samples from all vehicle-treated mice were pooled for sequencing, and samples from all ipatasertib-treated mice were pooled together. Gene expression data for the treated mice is available upon request.

#### OT

The data[26] comes from high-grade serous ovarian carcinoma (HGSOC) patients. We used batch 2, which includes four patients pooled together for sequencing. The dissociated cells were labelled with TotalSeq™-B anti-human hashtag antibodies.

#### EMBRYO

The EMBRYO data[25] is derived from embryonic day 18.5 mouse brain. All cells originate from a single mouse and were partitioned into 12 replicates using two reagents: MULTI-Seq lipid-modified oligos (LMO) and custom MULTI-Seq cholesterol-modified oligos (CMO). A high-throughput 96-well protocol was used to label the cells. Due to the use of an inbred mouse strain, SNP-based demultiplexing is not applicable.

#### PDX

For the PDX data[25], nuclei were extracted from a human ovarian carcinosarcoma patient-derived xenograft (PDX). Nuclei from four donors were then labelled using CellPlex oligos, custom MULTI-Seq cholesterol-modified oligos (CMO), and TotalSeq`™`-A anti-nuclear pore complex antibodies, respectively.

#### PBMC

In the PBMC data[1], peripheral blood mononuclear cells (PBMCs) from eight human donors were labelled with antibody oligos.

## Declarations

### Ethics approval and consent to participate

Not applicable

### Consent for publication

Not applicable

### Availability of data and materials

See the section of “Dataset”. The treated mouse data and benchmarking code are available at: https://zenodo.org/records/16976382. The source code of CMDdemux is available at: https://github.com/jiananwehi/CMDdemux.

### Authors’ contributions

JW developed the method and conducted all analyses. LC contributed to the statistical methods of CMDdemux and deMULTIplex2. DB and CC contributed to the interpretation of low-quality data. TS oversaw the entire project. All authors read and approved the final manuscript.

## Acknowledgements

We would like to thank George Howitt for providing the data. Our thanks also go to Chris Woodruff and Marie Trussart for their comments on the manuscript.

